# DDX24, a D-E-A-D box RNA helicase, is required for muscle fiber organization and anterior pole specification essential for head regeneration in planarians

**DOI:** 10.1101/2021.01.21.427618

**Authors:** Souradeep R. Sarkar, Vinay Kumar Dubey, Anusha Jahagirdar, Vairavan Lakshmanan, Mohamed Mohamed Haroon, Sai Sowndarya, Ramanathan Sowdhamini, Dasaradhi Palakodeti

## Abstract

Planarians have a remarkable ability to undergo whole-body regeneration. The timely establishment of polarity at the wound site followed by the specification of the organizing centers- the anterior pole and the posterior pole, are indispensable for successful regeneration. In planarians, polarity, pole, and positional-information determinants are predominantly expressed by muscles. The molecular toolkit that enables this functionality of planarian muscles however remains poorly understood. Here we report that SMED_DDX24, a D-E-A-D Box RNA helicase and the homolog of human DDX24, is critical for planarian head regeneration. DDX24 is enriched in muscles and its knockdown leads to defective muscle-fiber organization and failure to re-specify anterior pole/organizer. Overall, loss of DDX24 manifests into gross misregulation of many well-characterized positional-control genes and patterning-control genes, necessary for organogenesis and tissue positioning and tissue patterning. In addition, wound-induced Wnt signalling was also upregulated in *ddx24* RNAi animals. Canonical WNT-βCATENIN signalling is known to suppress head identity throughout bilateria, including planarians. Modulating this Wnt activity by *β-catenin-1* RNAi, the effector molecule of this pathway, partially rescues the *ddx24* RNAi phenotype, implying that a high Wnt environment in *ddx24* knockdown animals likely impedes their normal head regeneration. Furthermore, at a sub-cellular level, RNA helicases are known to regulate muscle mass and function by regulating their translational landscape. *ddx24* knockdown leads to the downregulation of large subunit ribosomal RNA and the 80S ribosome peak, implying its role in ribosome biogenesis and thereby influencing the translational output. This aspect seems to be an evolutionarily conserved role of DDX24. In summary, our work demonstrates the role of a D-E-A-D box RNA helicase in whole-body regeneration through muscle fiber organization, and pole and positional-information re-specification, likely mediated through translation regulation.

## INTRODUCTION

Planarian flatworms have exceptional ability to undergo whole-body regeneration (1, 2). Although this observation has fascinated scientists for centuries, the long-standing question of how this is accomplished remains to be completely understood. One critical determinant of planarian regeneration is the timely establishment of regeneration polarity at the wound site. Anterior facing wounds regenerate the head whereas posterior facing wounds regenerate the tail-this is known as regeneration polarity. Anterior polarity, for example, is determined by the expression of the Wnt antagonist *notum* at anterior facing wounds (3). This process is then followed by anterior-pole determination (4–9). The anterior-pole is the Spemann-Mangold organizer equivalent in planarians (10–12). Therefore, precise spatial and temporal determination of this pole is necessary for the head regeneration as well as patterning of different tissues within the head primordia. Further, in the intact animal, constitutive and regional expression of various morphogens and other positional-control genes (PCGs) is necessary for the maintenance of size, shape, and positional identity for different tissues (10,13–19). During regeneration, it becomes necessary to adjust the expression of these PCGs to reset positional information within the existing animal fragment (20–23). Factors regulating PCG expression, critical for the re-establishment of head-trunk-tail identity, remains an active area of investigation.

Planarian muscles, in addition to their canonical roles in locomotion and skeletal support, predominantly express all the positional-control genes and morphogens associated with positional information establishment and patterning (24, 25). Furthermore, muscles in planarians also act as ‘structural scaffolds’ for organ regeneration and perform the role of connective tissue by expressing many of the matrisome related genes (26–28). Therefore, given the multi-faceted functional role of muscle in regeneration, defect in muscle organization or loss of muscle associated factors inadvertently leads to defective regeneration (26,29–32). Molecules and mechanisms that underlie this functional versatility of planarian muscles however remain under-explored.

Post-transcriptional regulatory processes essential for fine-tuning gene expression have gained importance because of their critical role in various cellular functions (33–38). RNA helicases play a central role in the milieu of post-transcriptional gene regulation by modulating the secondary and tertiary structure of RNA, thereby regulating RNA\RNA and RNA\protein interaction (39, 40). Further, multiple lines of investigation from vertebrates show that D-E-A-D Box RNA helicases are essential for maintaining skeletal muscle mass, architecture, and function as well as muscle repair and regeneration via regulating ribosome biogenesis (41–43). Here, we report the function of DDX24, a D-E-A-D Box RNA helicase, in planarian head regeneration. *ddx24* was expressed in a subset of neoblasts, neoblasts primed for muscle fate, and also in the differentiated muscles. We custom generated a polyclonal antibody against DDX24 and found that the protein was particularly enriched in longitudinal and diagonal muscle fibers. Knockdown of *ddx24* resulted in muscle fiber disorganization and abnormal expression of anterior-pole markers like *zicA, foxD, and notum*, suggesting a defect in the anterior-pole specification. Loss of DDX24 also resulted in gross misregulation of many well-characterized patterning-control genes and positional-control genes, leading to defective regeneration. One such well-studied positional-control gene is *wnt1.* Although wound-induced expression of *wnt1* happens both in posterior and anterior blastema at early time-points of regeneration, this expression subsequently converges to the posterior tip by later stages of regeneration (44). However, in *ddx24* KD animals, Wnt activity is upregulated both in the posterior and anterior regenerating tissues compared to the controls. Modulating this Wnt activity by knocking down *β-catenin-1*, the effector molecule of this pathway, in the *ddx24* KD background, partially rescued the regeneration defect. Therefore, it is likely that DDX24 is required for regulating Wnt activity essential for proper localization of signaling centers essential for regeneration.

Like many other tissues, muscle function was shown to be critically dependent on its translational landscape, which in turn is primarily controlled by regulating ribosome biogenesis (42,45–49). Knockdown of *ddx24* in planarians resulted in reduced levels of large subunit ribosomal RNA (28S rRNA) as well as reduced 80S ribosome peak. This aspect appears to be an evolutionarily conserved role of DDX24 (50, 51). Since DDX24 protein was enriched in muscles, these results suggested that DDX24 could enable the functional state of these muscles by regulating their translational output via ribosome biogenesis.

## RESULTS

### DDX24 is necessary for anterior regeneration

D-E-A-D box helicases are highly conserved RNA binding proteins that play critical roles in numerous aspects of RNA metabolism. We performed an RNAi screen aimed at finding different RNA helicases essential for regeneration in planarians. Here we report that SMED-DDX24, the planarian homolog of human DDX24, was one of D-E-A-D Box RNA helicases necessary for anterior regeneration (Figure1 figure supplement 1a). While planarians regenerate all missing tissues within 7 days post-amputation (DPA), *ddx24* knockdown (KD) animals failed to do so. All the control animals regenerated their eyespots by 7 DPA whereas *ddx24* KD animals lacked the eyespots (Figure 1a). Photoreceptor in *Schmidtea mediterranea* is characterized by the presence of pigment cup and photoreceptor neurons (PRN). Immunostaining with anti-arrestin antibody and RNA-FISH for *opsin*, which cumulatively marks the PRN, optic chaisma, and cell body of the PRN (52, 53), showed defective photoreceptor organization in the *ddx24* KD animals (Figure 1b). We then sought to investigate the status of other organ systems in our knockdown animals. In planarians, the cephalic ganglia, i.e., its brain, is a bilobed structure (54), which can be visualized by the staining with nuclear dyes such as Hoechst or by immunostaining with anti-Gα q/11/14 antibody. In *ddx24* RNAi animals, the planarian brain either failed to regenerate completely, or a rudimentary brain was formed compared to the control animals (Figure 1c).

**Figure 1.**
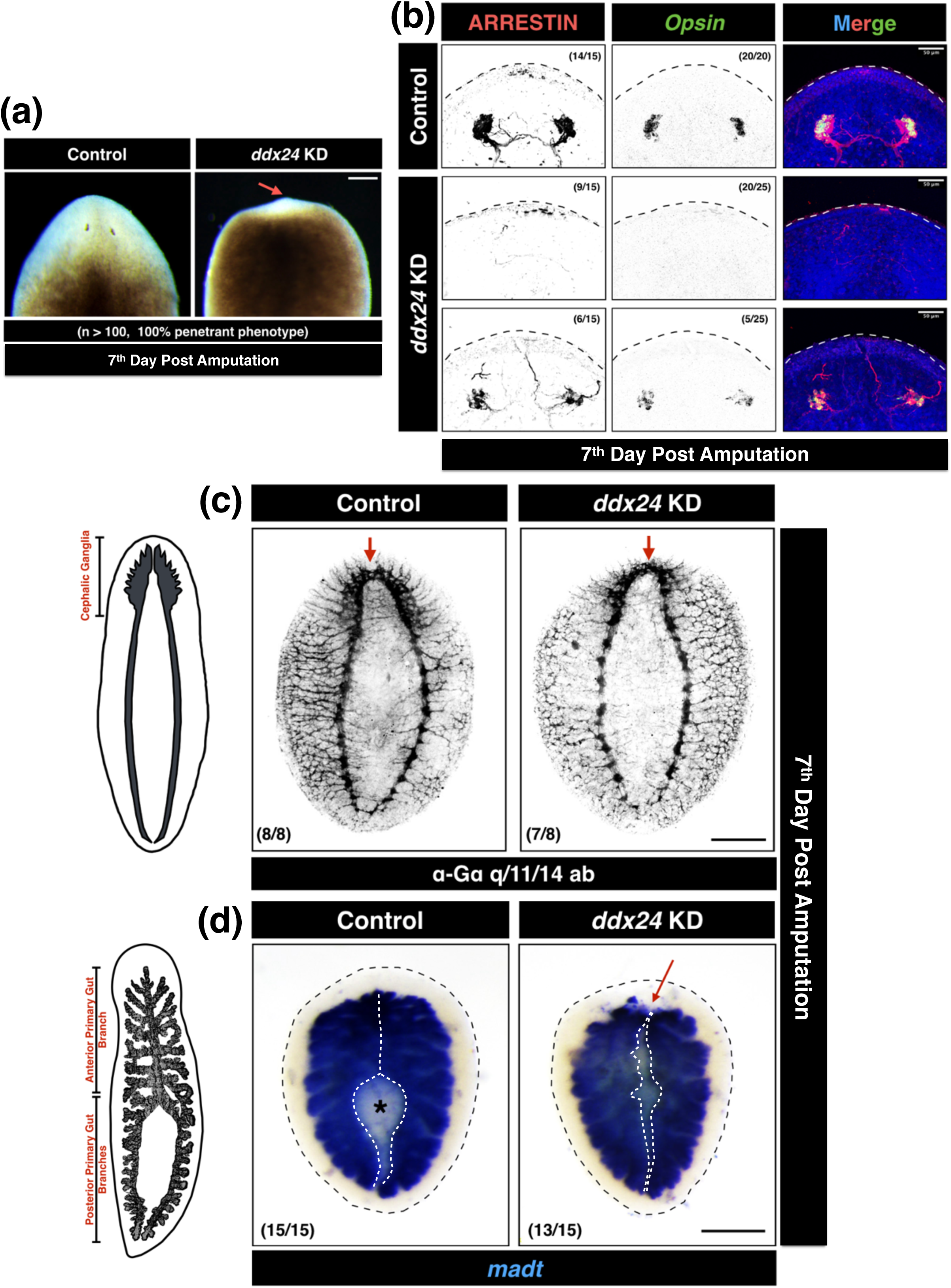
DDX24 is necessary for anterior regeneration in planarians. **(a)** Dark-field images of control and *ddx24* KD tails regenerating head at 7th-day post-amputation. Eyespots are lost in absence of DDX24 (red arrows) (n > 100, 100% penetrant phenotype. Scale bar: 200 µm) **(b)** Inability to regenerate eyes in *ddx24* KD corroborated by RNA-FISH for *Opsin* as well as immunostaining using anti-arrestin antibody. Either no eye structures were formed, or defective and rudimentary structures were formed. (Scale bar: 50 µm) **(c)** Immunofluorescence using anti-Gα-q/11/14 antibody revealed defective cephalic ganglia regeneration in *ddx24* KD (Scale bar: 200 µm). Animals mostly regenerated an underdeveloped rudimentary brain as shown here. In extreme cases, no cephalic ganglia were observed. These observations were in agreement with RNA insitu for *pc2* as well as immunostaining using an anti-synapsin antibody, both of which also mark the cephalic ganglia (data not shown). **(d)** RNA-insitu for *mat*, a bonafide gut marker. The anterior branch of the gut failed to regenerate in *ddx24* KD animals (red arrow). Also, no pharyngeal cavity (marked by asterisk *) was observed (Scale bar: 200 µm).

Planarian flatworms belong to the order tricladida characterized by the presence of three primary gut branches-one anterior primary gut branch that splits pre-pharyngeally into two posterior primary gut branches (55). dsRNA treated animals were amputated post-pharyngeally such that the posterior gut branches merge and then subsequently regenerate the entire anterior gut branch (Figure 1- figure supplement 1b). RNA in-situ for *mat*, a gut marker (56), clearly showed that *ddx24* KD animals failed to regenerate the anterior branch of the intestine (Figure 1d). Also, no pharyngeal cavity (marked by Asterix *) was observed in these knockdown animals. This was indicative of defective pharynx regeneration. Lack of pharynx in *ddx24* KD animals was also corroborated by 6G10 staining that marks pharyngeal muscles (Figure 1- figure supplement 1e) (57).

Together, our data shows that loss of DDX24 completely impaired anterior regeneration. Posterior/tail regeneration was also defective in absence of DDX24 (Figure 1- figure supplement 1f, g). Overall, DDX24 is necessary for regeneration in planarians.

### DDX24 protein is enriched in a subset of muscle fibers

Since we observed gross regeneration defects in *ddx24* knockdown, to better understand the localization and function of DDX24, we custom generated a polyclonal antibody against the peptide, SPKSLNADMQQKMRLKKLE, specific to planarian DDX24. Knockdown of *ddx24* followed by western blot and immunostaining with this antibody showed decreased levels of DDX24, thereby validating the efficacy and specificity of our antibody (Figure 2- figure supplement 2a-c). Immunostaining using this antibody prominently labeled body wall muscles (Figure 2a). Planarian body wall musculature (BWM) consists of three layers-circular, diagonal, and longitudinal muscle fibers (31). Our antibody stained the diagonal and longitudinal muscle fibers conspicuously. In contrast, circular muscle fibers showed a lower expression of DDX24 (Figure 2b). Overall, our immunofluorescence data showed that DDX24 was predominantly present in the longitudinal and diagonal muscle fibers compared to any other tissue in the animal. To further validate the enrichment of DDX24 in muscle, we employed double-RNA-FISH (dFISH) (58) to uncover the co-expression of *ddx24* with *collagen* and *bwm1*, bonafide pan-muscle markers in planarians (24,27,32). We found that a significant number of *ddx24+* cells were also positive for *collagen* (Figure 3a) and *bwm1* expression (Figure 3- figure supplement 3a). For instance, 65.8 ± 9.8% of *collagen+* cells and 63.4 ± 7.6% of *bwm1+* cells also expressed *ddx24.* This together with the immunofluorescence data demonstrated that DDX24 protein and its transcript are expressed in muscles. MyoD is the master transcription factor that specifies longitudinal muscle fibers in planarians (31). Since DDX24 protein was enriched in longitudinal muscles, we also validated this observation by performing double-RNA-FISH for *ddx24* and *myoD*. Co-localization experiments showed that 54.2 ± 0.7% of the *myoD+* cells also expressed *ddx24* (Figure 3b). NKX1.1 on the other hand specifies circular muscle fibers and DDX24 also had lower expression in these fibers compared to longitudinal or diagonal muscle fibers. In agreement with the latter observation, we found that a relatively lower percentage of *nkx1.1+* cells, i.e., 21.3 ± 7.8%, co-expressed *ddx24* (Figure 3- figure supplement 3b).

**Figure 2.**
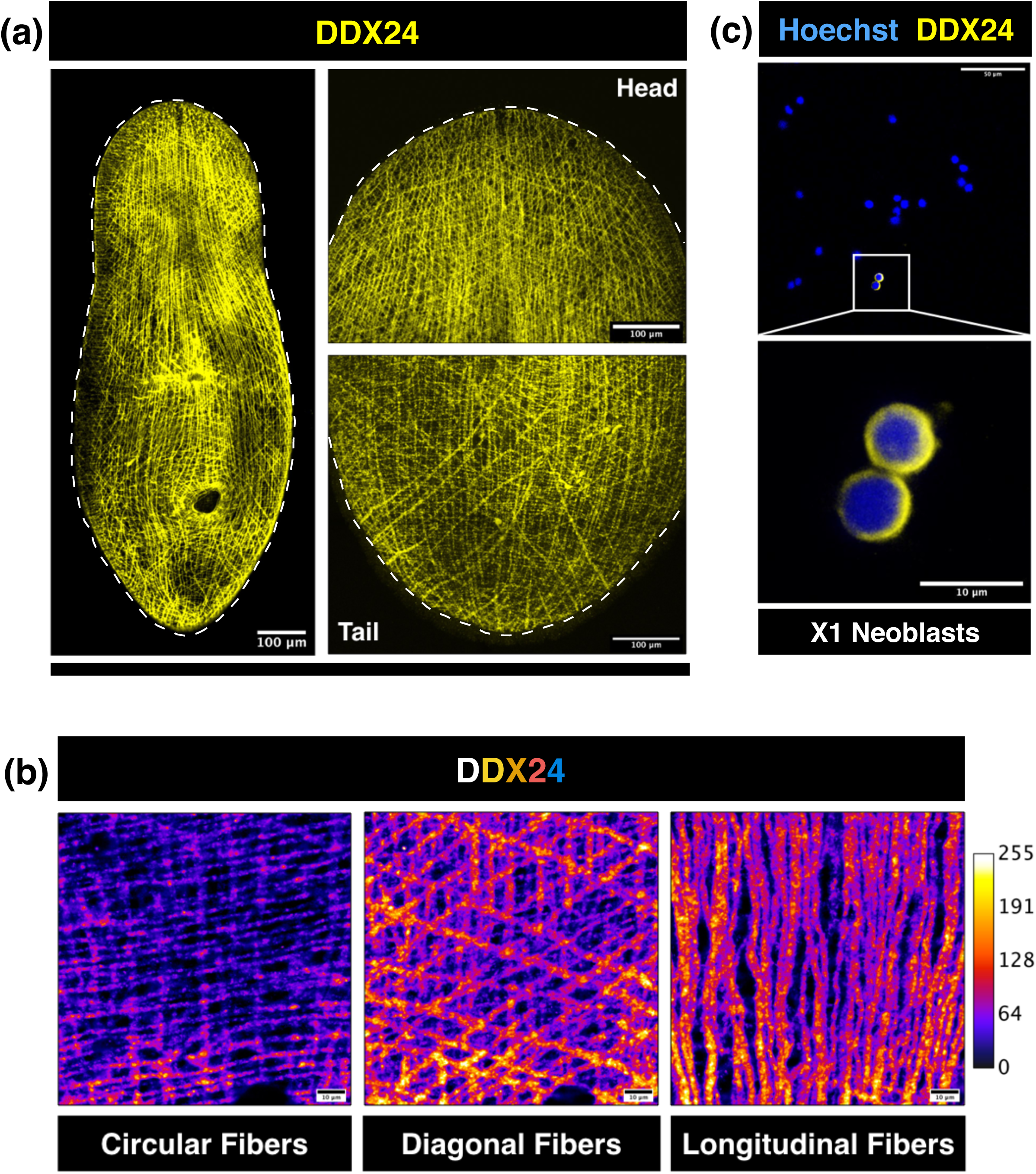
DDX24 protein was present in a subset of planarian body wall muscle fibers and X1 neoblasts. **(a)** DDX24 immunofluorescence in intact planaria. The ventral surface from head and tail regions zoomed-in for greater clarity. The fibrous pattern, as seen here, is characteristic of planarian body wall muscle fibers. (NOTE: All the animals here are different) (Scale bar: 100 µm, n > 20) **(b)** Amongst different muscle fiber subsets, DDX24 protein showed enriched expression in diagonal and longitudinal muscle fibers compared to circular muscle fiber. (Scale bar: 10 µm) **(c)** DDX24 protein was expressed in a subset FACS sorted X1 neoblasts. (Scale bar: 50 µm (top) and 10 µm (bottom)).

**Figure 3.**
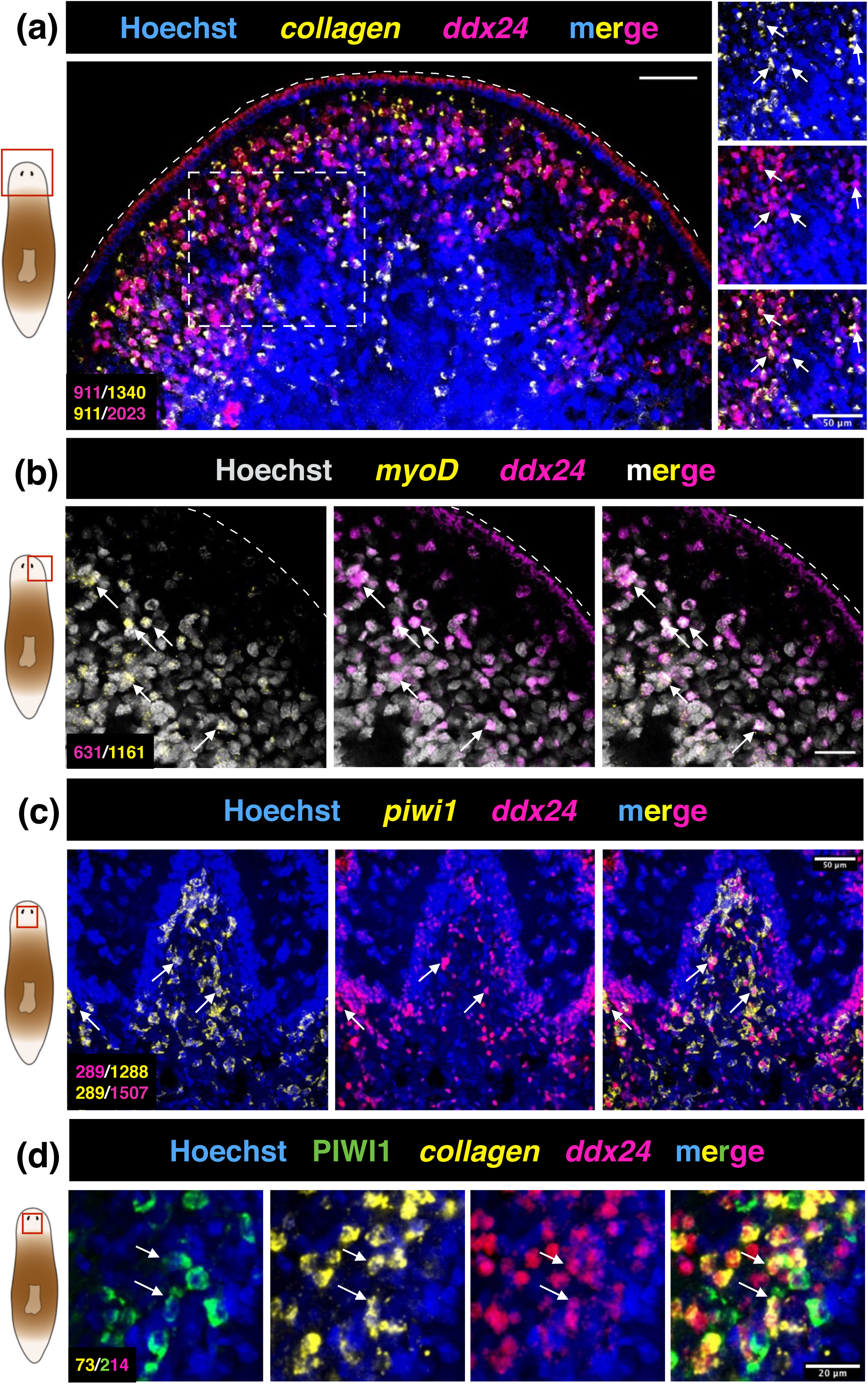
*ddx24* mRNA was expressed by a subset of muscles, neoblasts, and neoblasts primed for muscle fate. **(a)** Double-RNA-FISH confirmed that a subset of *ddx24* mRNA+ cells was also positive for *collagen*, a bonafide muscle marker. 65.8 ± 9.8% of *collagen*+ cells were also positive for *ddx24* whereas 46.7 ± 9.4% of *ddx24*+ cells were positive for *collagen*. (Scale bar: 50 µm) **(b)** DDX24 protein was found to be enriched in longitudinal muscle fibers. *myoD* is the master transcription factor specifying this muscle subtype in planarians. 54.2 ± 10.7% of *myoD*+ cells also co-expressed *ddx24*. (Scale bar: 50 µm) **(c)** A subset of *ddx24*+ cells are also *piwi1*+ neoblasts. 22.1 ± 4.6% of *piwi1*+ cells were also positive for *ddx24* whereas 18.8 ± 3.4% of *ddx24*+ cells were positive for *piwi1*. (Scale bar: 50µm) **(d)** *ddx24* is expressed in a class of stem cells primed for muscle fate. We found that 36.5 ± 9.7% of PIWI1+ *ddx24*+ double-positive cells were positive for *collagen* too. (Scale bar: 20 µm) (Note: The actual number of cells counted in each case is mentioned within the figure, with colour code. Every quantification percentage mentioned here was calculated in the entire volume specified by the red box on the adjacent planarian cartoon. 6 animals were used for quantification, except in (d) where 4 animals were used)

Previously published transcriptome data (59) revealed the expression of *ddx24* in planarian stem cells i.e., neoblasts. Immunostaining for DDX24 on FACS sorted X1 cells (proliferating neoblasts) showed the presence of DDX24 in a subset of these cells (Figure 2c). This observation was further supported by the double-RNA-FISH, which showed that 22.1 ± 4.6% of *piwi-1+* cells, a pan-neoblast marker, also expressed *ddx24* (Figure 3c). Since we found DDX24 protein in muscles and stem cells, we hypothesized that a fraction of the neoblasts that expressed the *ddx24* could be a neoblast population primed for muscle lineage. To test this, we performed double-RNA-FISH using *in situ* probes for *ddx24* and *collagen*, in combination with immunostaining using an antibody against PIWI-1 protein, a marker for pluripotent neoblasts and early progenitors (i.e., immediate neoblast progeny) (60). Our results showed that 36.5 ± 9.7% of *ddx24*-PIWI1 double-positive cells were also positive for *collagen* (Figure 3d). This suggested that a subset of *ddx24*+ neoblasts were indeed primed for muscle fate. This finding was further corroborated by multiple single-cell RNA sequencing datasets (61, 62) which also show the expression of *ddx24* in the muscle primed neoblast population (Figure 3- figure supplementary 3e-g).

It is worth mentioning that even though single-cell RNA-seq datasets point towards the expression of *ddx24* transcript in other lineages, like the epidermis, gut, neural, and pharynx, our antibody did not detect DDX24 protein in any of these differentiated planarian tissues. However, in intact animals, DDX24 expression was detected in cells other than muscle fibers whose identity we were unable to determine (Figure 2- figure supplement 2d).

### Loss of DDX24 leads to defective muscle fiber integrity and organization

Since DDX24 protein was enriched in muscles, we next investigated the status of muscle fibers in *ddx24* KD animals. Body wall muscle fibers run across the entire body wall column in planarians. Each fiber is continuous and these are of three kinds-circular, diagonal, and longitudinal (25, 31). Loss of DDX24 led to a severe defect in muscle fiber organization and integrity (Figure 4 and Figure 4- figure supplement 4d). These indentations and fractures in muscle fiber architecture were predominately localized around the central region on both the dorsal as well as on the ventral surface of the animal (Figure 4- figure supplement 4c video). Dark-field images and Hoechst staining revealed that there was no overall indentation or lesions on the surface of these animals (Figure 4 and Figure 1- figure supplement 1f) suggesting that this specific phenotype was restricted to the muscle compartment. Albeit at a lower frequency, we also observed these muscle fractures at the anterior tip of *ddx24* KD animals (Figure 4- figure supplement 4a). This was interesting because this region houses specialized muscle cells known as anterior pole cells which acts as the head-organizer essential for head regeneration (4,7–9). Further, we observed that the loss of DDX24 did not affect the expression of *collagen* suggesting that DDX24 was not necessary for *piwi1+* neoblasts to differentiate into *collagen+* muscles (Figure 4- figure supplement 4b). *myoD* levels too remained unchanged in absence of DDX24 (Figure 4- figure supplement 4j). On similar lines, even though DDX24 was also expressed by a population of neoblasts, we found that loss of DDX24 did not affect their maintenance, proliferation, or their fate commitment to different lineages (Figure 4- figure supplement 4e-i). Together, our data suggest that DDX24 predominantly functions at the level of muscle fiber organization rather than affecting gross muscle fate specification.

**Figure 4.**
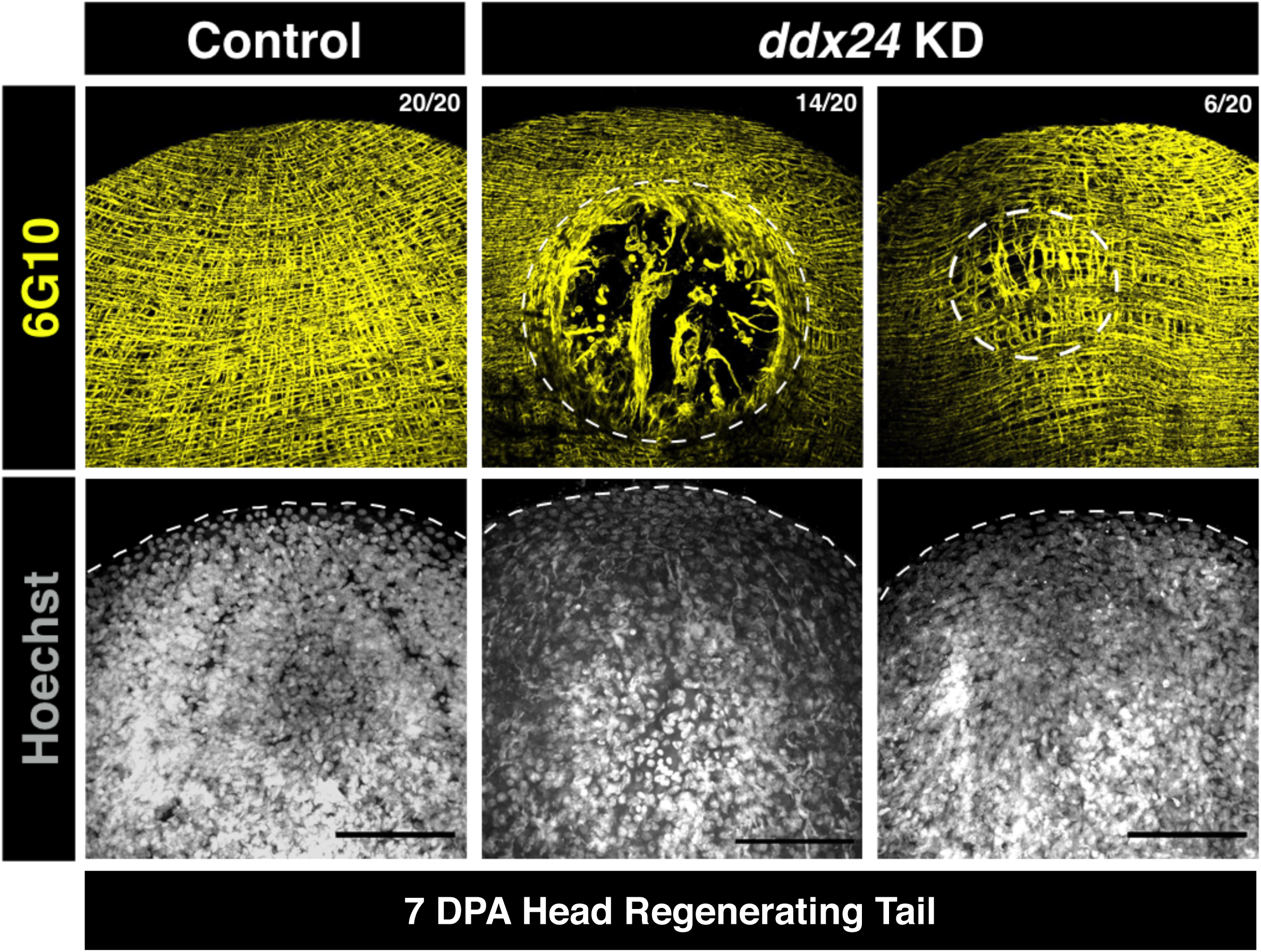
Loss of DDX24 leads to a defect in muscle fiber organization and integrity. 6G10 immunostaining revealed the loss of muscle fiber organization and their integrity in *ddx24* RNAi animals. Indentation and fracture in muscle fiber organization were observed on both the dorsal as-well-as on the ventral surface. Dark-field images and Hoechst staining revealed that there was otherwise no overall indentation or hole or lesion on these animals. This novel phenotype was restricted only to the muscle compartment. (Scale bar: 50 µm) (7- day post regenerating tail fragments)

### DDX24 is necessary for anterior pole re-specification during regeneration

Head regeneration and patterning of different tissues within the head primordia is critically dependent on the spatial and temporal determination of the anterior pole. The anterior pole is marked by the combined expression of transcripts like *zicA*, *foxD*, *notum*, in a neoblast progeny committed for muscle lineage (7–9). This fate commitment occurs between 24 to 72 hours post-amputation (hpa) exclusively at anterior facing wounds. RNA-FISH for *ddx24* showed its expression in the anterior blastema by 72 hpa. Interestingly, we noted co-expression of *ddx24* with *zicA+ foxD+* anterior pole progenitors at this time point of regeneration (Figure 5- figure supplement 5a). We further noted co-expression of *ddx24* with *zicA+ foxD+* anterior pole in intact animals (Figure 5a). Multiple single-cell RNA sequencing datasets further corroborated this observation that *ddx24* was indeed co-expressed by anterior pole cells- the expression of *ddx24* in the NB6 cluster (62) (Figure 3- figure supplement 3f) as well as co-expression of genes like *zicA, fst, prep, pbx, notum, sFPR-1, foxD, and nr4* in *ddx24+* single cells from Fincher et al., 2018 (61) (Figure 3- figure supplement 3g).

**Figure 5.**
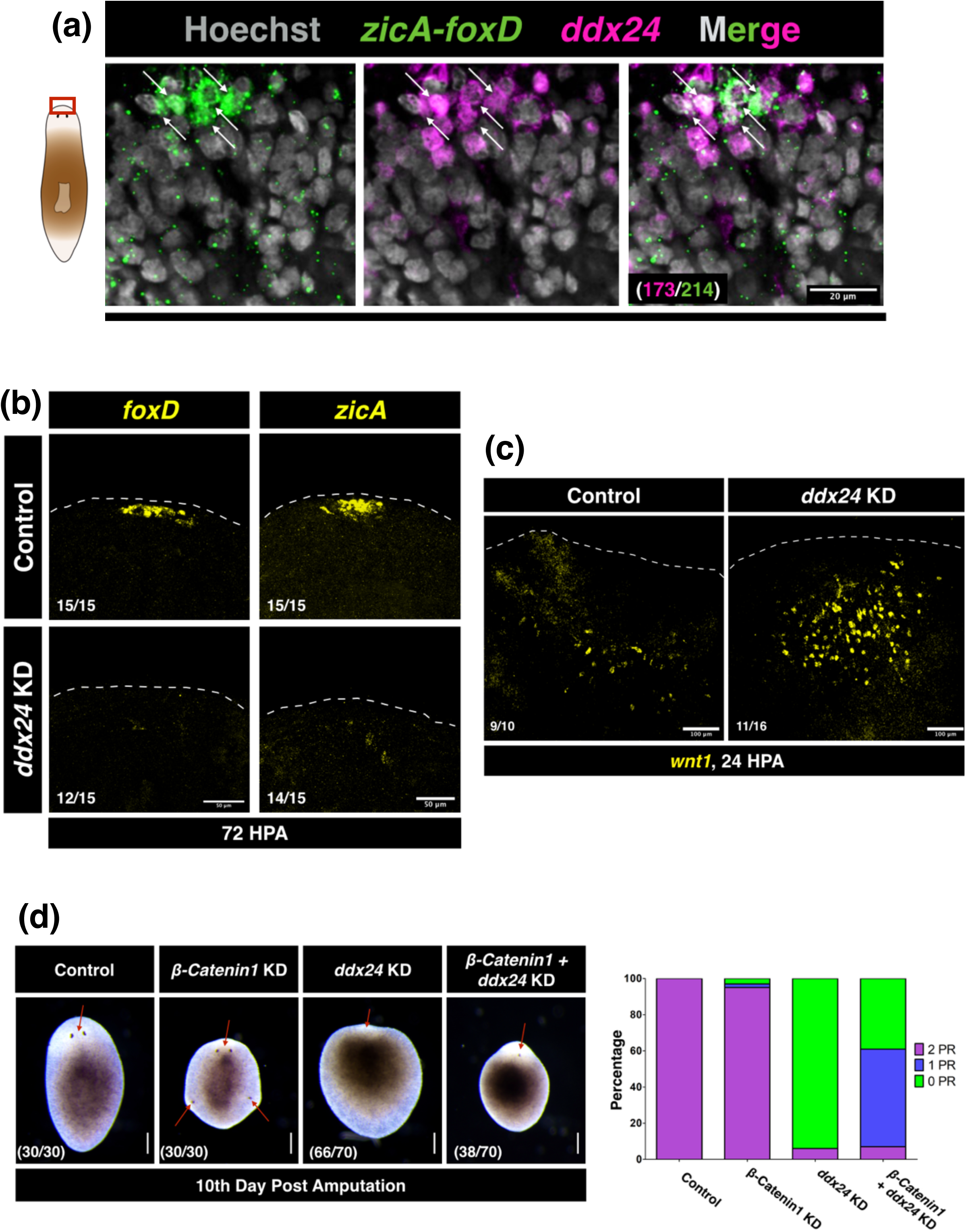
Anterior pole cells express *ddx24* and the loss of DDX24 leads to defective anterior pole re-establishment during regeneration. **(a)** The anterior pole (marked here by *zicA-foxD* expression), are specialized *collagen*+ muscle cells that are essential for head regeneration and patterning of anterior tissues. 79.9± 9.5% of *zicA-foxD*+ cells in intact planarians co-expressed *ddx24* at the head tip. (Scale bar: 20 µm, counting performed on 6 animals) **(b)** Loss of DDX24 eliminated *foxD* and *zicA* expression at the head tip in 72 hpa tail fragments regenerating head. **(c)** By 24 hpa, wound-induced *wnt1* expression reduces at the anterior blastema in control animals whereas higher number *wnt1+* cells are still observed in *ddx24* KD animals. (Scale bar: 100 µm) **(d)** Partial rescue of the eye-less *ddx24* RNAi phenotype by *β-catenin-1* RNAi in tail-fragments regenerating head (10-day post amputation animals). (Scale bar: 200 µm)

It is well established that loss of pole determinants like *zicA, foxD*, and *notum* leads to a failure in head regeneration (3,7–9). *ddx24* RNAi animals also failed to specify *zicA+ foxD+* pole cells by 72 hpa at the anterior tip (Figure 5b). Intriguingly, at the same timepoint, *ddx24* KD animals either failed to completely specify *notum+* anterior pole or showed gross mis-patterning of *notum* expression (Figure 5- figure supplement 5b). Next, we performed bulk RNA sequencing from *ddx24* KD and control animals to identify different genes and pathways mis-regulated after *ddx24* RNAi. We found that genes necessary for pole specification cum head regeneration like *pbx/extradenticle*, *follistatin* (*fst)*, and *foxD* (5–7,9,63,64) were down-regulated *ddx24* knockdown animals (Figure 5- figure supplement 5d). We also compared our sequencing data with another recently published anterior pole enriched transcriptome dataset (65). This enabled us to identify two distinct cohorts of anterior pole enriched transcripts- one cohort was downregulated whereas the other was upregulated in absence of DDX24 (Figure 5- figure supplement 5e). These data, therefore, suggest that the genetic program associated with anterior pole specification during head regeneration is abnormal in *ddx24* knockdown animals. Defect in muscle fiber organization and pole specification suggested that positional-information re-specification was likely defective in absence of DDX24 (5–9,31). This is important because the expression of positional-control genes need re-wiring within the existing animal fragment during regeneration (23). This allows for re-specification of head-trunk-tail identities necessary for organogenesis, tissue-positioning, and tissue-patterning to proceed in a bonafide manner (13–15,66). Re-analyzing previously published single-cell RNA sequencing data (61) suggested that a sizable population of *ddx24+* cells co-expressed many of these positional-control genes and patterning genes (Figure 3- figure supplement 3g). We then performed RNA-FISH for some of these genes like *ndl-3, ndl-5, sFRP-1, sFRP-2,* and *slit-1*, and found that their expression was mis-regulated in absence of DDX24 (Figure 5- figure supplement 5c). This finding was again corroborated by our bulk RNA-sequencing analysis that indeed many well-characterized positional-control genes and patterning genes such as *netrin-3*, *ndl-2, ndk, sFRP-1, slit-1, fz-4-2, fz-5/8-3,* and, *wnt-11-1* (16,22,23,67,68) were downregulated *ddx24* KD animals (Figure 5- figure supplement 5d). These observations thus indicate that DDX24 is required for the proper expression of positional control genes necessary for proper head regeneration and patterning in planarians.

### DDX24 suppresses would-induced expression of *wnt1*

Canonical WNT-βCATENIN signaling is required for tail specification with concomitant suppression of head identity throughout bilateria, including planarians (69–72). Failure of *ddx24* KD planarians to regenerate head hinted at the possibility of aberrant Wnt signaling in these animals. Although expression of *wnt1*, a canonical Wnt signaling ligand, at the wound site is a generic response to any wounding in planarians (44, 73), head-regenerating wounds however completely suppress the expression of *wnt1* at the wound site by 24 to 30 hpa. Suppression of this *wnt1* expression at head-regenerating wounds is necessary for neoblasts to specify *zicA+ notum+* anterior-pole cells, in absence of which head regeneration fails (8, 64). Upon RNA-FISH, we found that the number of wound-induced *wnt1*+ cells was greatly increased in absence of DDX24 (Figure 5c and Figure 5- figure supplement 5f). In addition, our transcriptome data also revealed that the expression of *wnt-11-4/wntP3/wnt-4* (22, 23), another canonical Wnt signaling ligand, was significantly upregulated in *ddx24* KD animals (fold-change (*ddx24* KD/Control KD) = 3.14; adjusted p-value = 0.0009). Therefore, it is likely that the activity of canonical WNT-βCATENIN signaling is upregulated in absence of DDX24 and this could be one potential reason why *ddx24* KD planarians fail to regenerate their head. To further test this idea, we performed a double knockdown of *β-catenin-1* and *ddx24*. We decided to knockdown *β-catenin-1* because canonical Wnt ligands, WNT1 and WNT-11-4, both act via β-CATENIN-1 (22), a common effector of the canonical Wnt pathway. Approximately 54% of *β-catenin-1 ddx24* double knockdown animals regenerated at least one eyespot by the 10^th^ day of regeneration whereas less than 5% of *ddx24* single knockdown animals did so (Figure 5d). Therefore, this partial rescue of *ddx24* RNAi phenotype by *β-catenin-1* RNAi supports the hypothesis that higher Wnt activity in *ddx24* KD animals is one potential reason why these animals fail to regenerate their head. This data is in alignment with previously published results that upregulation of Wnt activity (either through *fst* RNAi or *APC* RNAi) leads to defective head regeneration since a high Wnt environment leads to loss of *zicA+ notum+* anterior-pole cells (8, 64), something we observe in *ddx24* KD animals as well.

### *ddx24* KD leads to downregulation of ribosomal RNA processing machinery

We next sought to uncover one potential sub-cellular mechanism how DDX24, an RNA helicase, could affect muscle architecture and function, thereby affecting whole-body regeneration. GO analysis of our bulk RNA sequencing data revealed that the expression of many genes associated with ribosomal RNA processing machinery was de-regulated in *ddx24* KD animals (Figure 6a). Mak5, the yeast homolog of DDX24, was shown to be essential for large subunit ribosomal RNA maturation (50,51,74). In planarians, the qRT-PCR analysis revealed ∼40% downregulation in 28S rRNA levels upon *ddx24* KD whereas 18S rRNA levels remain nearly unchanged (Figure 6b). Therefore, our data hinted towards potential mis-regulation in translation mediated by reduced levels of mature 80S ribosomal. This was tested by performing polysome profiling on planarian lysates from control and knockdown animals.

**Figure 6:**
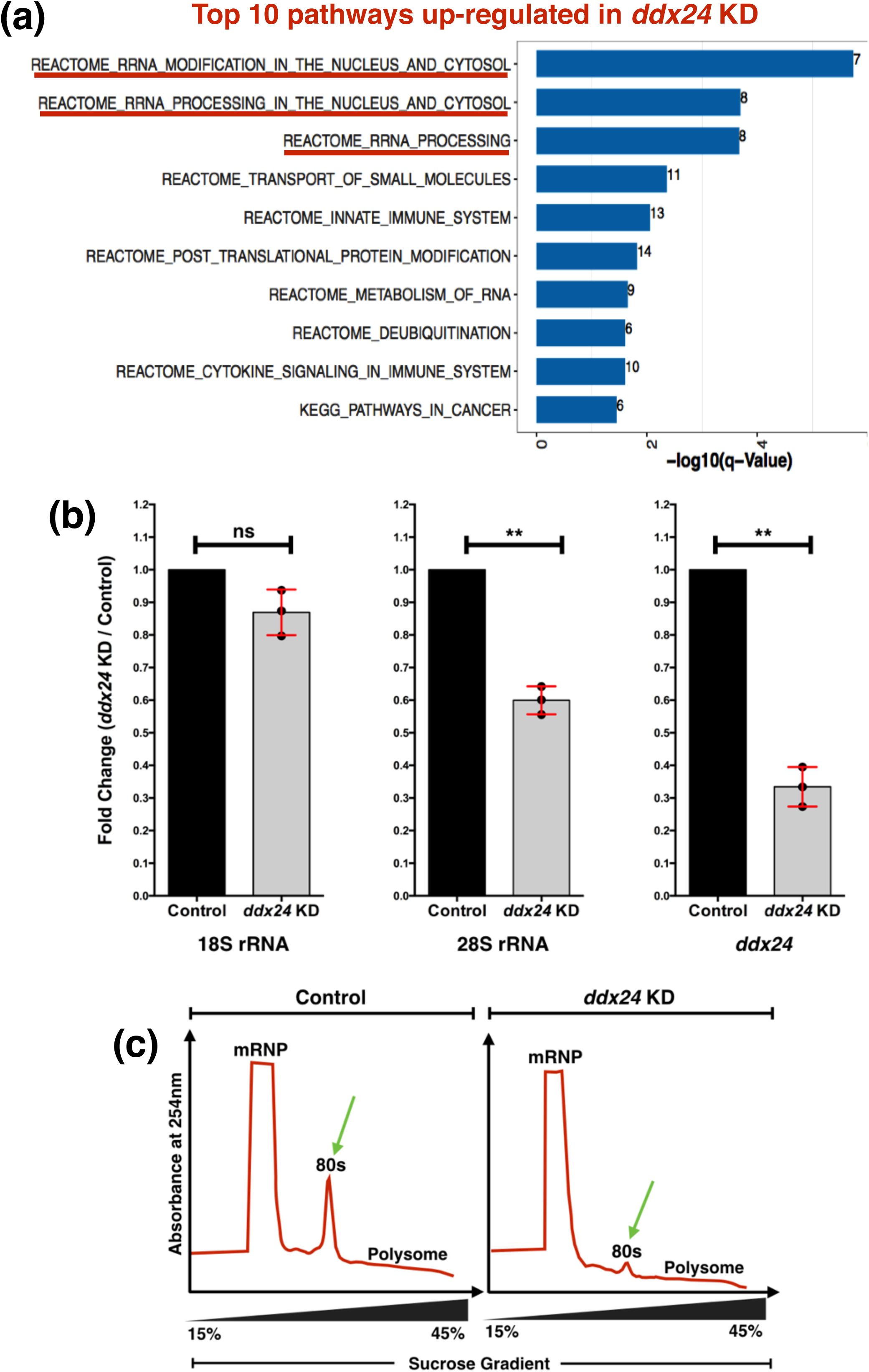
Loss of DDX24 leads to defective 28S ribosomal RNA processing and ribosome assembly. **(a)** Gene ontology for pathways and processes up-regulated in d*dx24* KD indicated that the expression of many genes associated with rRNA processing was affected in *ddx24* knockdown. **(b)** qRT-PCR for 18S rRNA, 28S rRNA, and *ddx24* transcript levels from 3DPA tail fragments from control and *ddx24* KD animals. All experiments were performed thrice. Mean ± SD indicated. Unpaired t-test was used to calculate statistical significance. Actin was used for normalization. (** implies p-value < 0.01) **(c)** Polysome profiling on 3DPA tail fragments from control and *ddx24* KD animals. In absence of DDX24, a reduction in the 80S ribosome peak was observed.

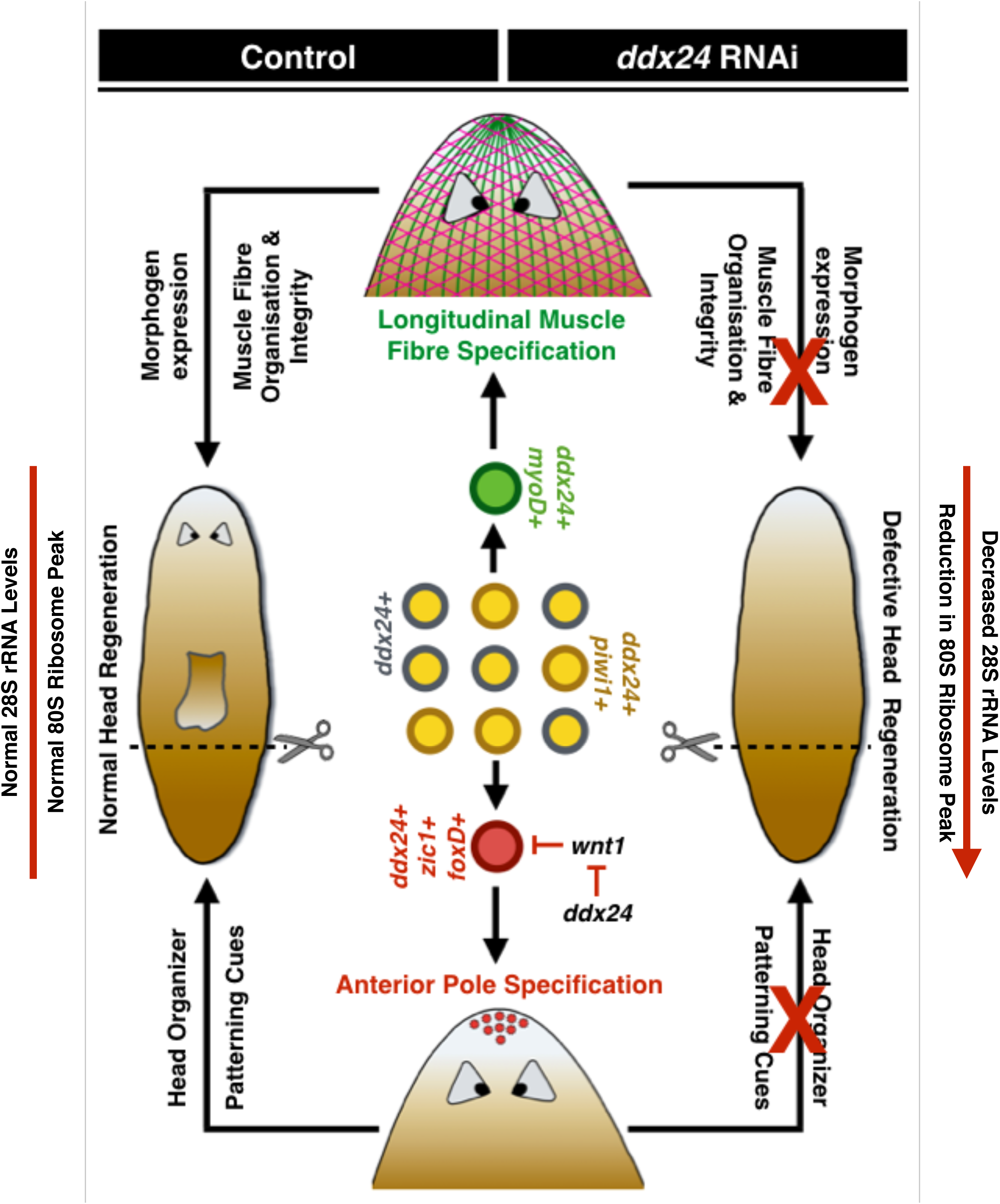

As predicted, the 80S ribosome peak was reduced in the absence of DDX24, which is suggestive of defective ribosome assembly leading to a perturbed translational landscape (Figure 6c). Since DDX24 protein was enriched in body wall muscles, and a subset of X1 neoblasts, we speculated that defective ribosome biogenesis in *ddx24* KD animals was predominantly restricted to these specific compartments. Therefore, it is likely that muscle defects in *ddx24* KD animals, both in terms of muscle fiber organization as-well-as failure to specify anterior-pole cells, could be a result of defective translational machinery in these cells. In conclusion, our work highlights the role of translation regulation mediated by a D-E-A-D Box RNA helicase in whole-body regeneration via maintaining muscle form and function.

## DISCUSSION

Planarians have a remarkable ability to undergo whole-body regeneration. Any part of their body lost to injury or amputation is functionally restored in a matter of weeks! It has therefore been a long-standing interest to understand the factors and mechanisms that enable this process. For example, successful polarity determination at the wound site is necessary for proper regeneration. The expression of a Wnt pathway antagonist *notum,* for instance, determines anterior polarity at an anterior-facing wound site (3). This process is followed by anterior pole re-specification. Anterior pole cells are marked by the co-expression of markers like- *notum, fst, foxD,* and *zicA* in neoblast progeny primed for muscle lineage (7–9). These clusters of cells are necessary for initiating head regeneration as well as positioning and patterning of different anterior tissues.

In planarians, muscles predominantly express all the pole determinants as well as other positional-control genes that are associated with head-trunk-tail identity re-specification within the regenerating animal fragment (24). Muscles also serve as structural scaffolds for organ regeneration (26). In vertebrates, connective tissues, like fibroblasts, migrate to the wound site, generate different tissue types as well as display positional memory-processes essential for limb regeneration (75–82). Interestingly, in planarians, muscles have additional connective tissue type roles too. They express all the matrisome related genes-these ECM glycoproteins, *perlecan*, *hemicentin-1,* for example, are necessary for maintaining spatial positioning of different mesenchymal cell-types in these flatworms (27, 28). However, the factors that underlie this tremendous functional versatility of planarian muscles remains poorly understood.

Previous studies in vertebrates have shown that regulating the translational landscape was necessary for maintaining the functional state, biomass, and architecture of muscles (42,45– 49). Translation can be regulated by different post-transcriptional processes such as ribosome assembly, mRNA stability, and regulation of translation initiation and elongation (83–86). D-E-A-D box (DDX) helicases, a highly conserved family of RNA binding proteins and well-known regulators of different post-transcriptional and translational processes, were shown to be critical for myogenic fate and function (41–43,87,88). For example, in zebrafish, DDX27 is necessary for the proliferation and differentiation of myogenic progenitor cells to myofibers as well as for muscle repair and regeneration post-injury. DDX27 is essential for rRNA synthesis and ribosome assembly in these cells and *ddx27* knockout downregulates the translation of transcripts necessary for myogenesis (42).

We conducted an RNAi screen for different post-transcriptional gene regulators, including members from the D-E-A-D box RNA helicase family, in *Schmidtea mediterranea*, the go-to model system to study whole-body regeneration. Here, we report that SMED_DDX24, the *Schmidtea mediterranea* homolog of human DDX24, is essential for its regeneration. Although DDX24 is conserved across eukaryotes, little is known about the function of this protein in regulating various biological processes. In this study, we have mostly focused our effort to understand how DDX24 governs head regeneration in planarians. We found that *ddx24* mRNA is expressed by a population of neoblasts, neoblasts primed for muscle fate, and differentiated muscles. However, the protein is highly enriched in body-wall muscle fiber, particularly diagonal and longitudinal muscle fibers. We also detected the protein in a subset of neoblasts. Although, the loss of DDX24 did not affect *piwi1+* neoblasts, their maintenance, proliferation, or their overt differentiation. However, the loss of DDX24 did severely affect muscle fiber organization and integrity. This defect was observed both on the dorsal and the ventral surfaces and with a lower frequency, on the tip of the regenerating head. Although *ddx24* was expressed in a subset of stem cells primed for muscle fate, our transcriptome and RNA-FISH data suggested that *ddx24* RNAi does not affect muscle lineage specification as such. Therefore, it is likely that DDX24 is required for organizing muscle fibers and not for their overall fate determination.

We also found the co-expression of *ddx24* with the anterior pole markers, both in intact and in 72-hour post-amputation head-regenerating animals. Besides, the loss of DDX24 eliminated these pole cells too. This raised the possibility that DDX24 was required for anterior pole determination. Moreover, *ddx24* KD animals had highly upregulated and ectopic expression of wound-induced *wnt1*. This was interesting since a high level of Wnt activity is known to inhibit the specification of *zicA+* anterior pole cells, thereby leading to defective head regeneration (8). Therefore, we modulated this ectopic Wnt signalling through *β-catenin-1* RNAi, the effector molecule of this pathway. This led to the partial rescue of the eyeless phenotype in *ddx24* RNAi animals. Hence, it is likely that the defective head regeneration in *ddx24* KD animals is due to their inability to specify *zicA+* anterior pole cells caused by higher levels of wound-induced Wnt activity. A similar mechanism of action was also shown in *follistatin (fst)* knockdown (66). Over induction of wound-induced *wnt1+* cells in *fst* RNAi animals impedes the formation of *notum+* anterior pole, thereby leading to defective head-regeneration. Besides, *wnt1* or *β-catenin-1* RNAi rectified *fst* RNAi phenotype as well. Interestingly, our data also showed reduced levels of *fst* and *notum* in *ddx24* KD. Although the specification of *foxD+* anterior-pole cells is independent of WNT-βCATENIN signalling (7), elimination of *zicA* somehow eliminates *foxD* expression by 72-hour post-amputation (9). Therefore, this could be one possible explanation for why *ddx24* RNAi animals also failed to specify *foxD+* pole cells. Furthermore, tail regeneration was also defective in absence of DDX24. Specification of *wnt1+* posterior-pole is necessary for tail regeneration (44, 68) and *ddx24* RNAi animals also had a very high level of ectopic *wnt1* expression at the tail-blastema. Although how this process unfolds is not understood, nevertheless it is very interesting that an abnormally high level of Wnt signalling could impede tail regeneration. Together, our data suggest that DDX24 plays an important role in modulating the levels of wound-induced Wnt activity, necessary for proper pole organization during regeneration.

The expression of many well-characterized positional-control genes was grossly mis-regulated in *ddx24* knockdown animals. Many of these genes are essential for tissue-positioning and tissue-patterning during regeneration, necessary for proper organogenesis. For example, *ndk* levels were downregulated in absence of DDX24. *ndk*, an FGF receptor-like gene (86), regulates cephalic ganglia regeneration by delineating its posterior boundary. Another example would be that of *slit-1*, an evolutionarily conserved gene necessary for proper medial-lateral positioning of regenerating organs (67), whose levels were also reduced in *ddx24* knockdown. It is worth recapitulating here that- (i) all these positional-control genes are predominantly expressed by muscles and (ii) loss of muscle fibers (for example *myoD* RNAi (31)) or loss of anterior pole determinants (7–9) results in abnormal positional-control gene expression. Therefore, to summarize, our study shows a novel role of DDX24 in regulating muscle fiber organization, anterior pole determination, and positional-information re-specification thereby enabling head regeneration in planarians.

Mak5, the yeast homolog of DDX24 is necessary for large subunit ribosome RNA biogenesis, in effect ribosome biogenesis (50,51,74). In planarians, we investigated the potential role of DDX24 on 28S rRNA levels. Although 18S rRNA levels remained unperturbed, the knockdown of *ddx24* led to a decrease in levels of 28S rRNA which subsequently led to a reduction in the 80S monosome peak. DDX24 knockdown, therefore, alters the translational landscape in planarians. In vertebrates, it has been shown that regulating the translatome is important for maintaining the fate and function of skeletal muscles. On similar lines, our study highlights the possibility that DDX24 regulates ribosomal biogenesis in planarian muscle cells via regulating 28S rRNA levels. This regulation in turn could be important for muscle function. However, the mechanism by which DDX24 regulates ribosome biogenesis remains to be investigated. D-E-A-D box RNA helicases are versatile proteins- they can directly interact with proteins as well as different species of RNA to elicit their specific function in a context-dependent fashion. Future interactome studies of DDX24 would help to understand precise molecular underpinnings through which it regulates whole-body regeneration in planarians.

## MATERIAL & METHODS

### Animal husbandry

The sexual strain of *Schmidtea mediterran*e*a* was used in this study. All animals were kept in 1X planaria media (1.6 mM NaCl, 1.0 mM CaCl2, 1.0 mM MgSO4, 0.1 mM MgCl2, 0.1 mM KCl, and 1.2 mM NaHCO3 pH adjusted to 7.0) in 20 degrees. Planarians were starved for at least one week before any experiment.

### RNA Interference

Sequence of *ddx24* (transcript id: dd_v6_Smed_7310_0_1) can be found on PlanMine (http://planmine.mpi-cbg.de) (59). Primer pair used to amplify *ddx24*: forward primer-5’-TGCCAACAGCAAGAACTCAG-3’; reverse primer-5’GACGGTGCGTTTTTCTTCTC-3’.

dsRNA synthesis and feeding were performed as described by Rouhana & colleagues (89). dsRNA against the green fluorescent protein (GFP) was used as RNAi control. Animals were fed three dsRNA thrice with a gap of three days after each feed. On the fourth day post last feed planarians were amputated post pharyngeal to generate tail fragments. 1.5µg of dsRNA per 10 µL beef-extract was used to make the food-dsRNA mix. qPCR primer pair used to check *ddx24* knockdown: forward primer-5’-GGATTTTTGAAACCGACACA-3’; reverse primer 5’-CCCAAATGCTAATGTTTTGC-3’. Our transcriptome data shows that *ddx24* KD leads to the downregulation of *ddx24’s* expression alone without affecting the transcript levels of other DEAD box RNA helicases which also shows the specificity of the dsRNA used for the study (Figure1 figure supplement 1 c,d)

### RNA insitu hybridisation and immunostaining

RNA-insitu hybridisation was performed as described by King and Newmark (58). RNA-FISH for *ddx24* was always performed using a cocktail of two non-overlapping fluorescein-labeled riboprobes. The primer pair used to generate the first set of riboprobe is- forward: 5’-TTGCGTGGAATGACGATGAG-3’; reverse: 5’-AATGTGGCCGAAAACACGAG-3’, and primer pair used to generate the second set of riboprobe is- forward: 5’-TGGCAGCAAGAGGTTTGGAT-3’; reverse: 5’-GTGGCCAGAAAGAGGGGACT-3’. The final concentration of riboprobes used was 1ng/ µL. The specificity of the riboprobes against *ddx24* mRNA was validated by performing RNA FISH one week after *ddx24* RNAi (Figure 3 figure supplement 3 c, d)

Immunostaining for Arrestin (a kind gift from Prof. Kiyokazu Agata), 6G10, SMED-PIWI1, and H3P was performed as described by previous papers from our lab (90–92). Anti-Gα q/11/14 (G-7) (1:200 Santa Cruz Biotechnology, sc-365906) antibody staining was performed in formaldehyde-fixed samples after RNA-FISH was performed on them. Fluorophore conjugated anti-mouse secondary antibody (Life Technologies) was used at a dilution of 1:1000, for two hours at room temperature.

For anti-DDX24 antibody immunofluorescence, planarians were sacrificed in 5% N-Acetyl Cysteine in 1X PBS for 7 minutes, rinsed 2X with 1X PBS, and fixed in ice-cold meth-Carnoy solution (methanol: chloroform: acetic acid in 6:3:1 ratio by volume) for 45 minutes in cold-room with mild nutation post which they were washed multiple times with cold methanol and eventually brought to room temperature. Animals were gradually rehydrated in graded methanol-0.5% PBSTx. 6% H2O2 (either in 1X PBS or 100% methanol. Muscle staining is better with methanol bleach whereas detection of the “other” cell types happens better with PBS bleach) under bright light was used to bleach animals. Rabbit anti-DDX24 antibody was used at 1:50 dilution and incubated at 4 degrees for 24 hours. Anti-Rabbit-HRP (1:1000, Cell Signaling Technology 7074) was used as the secondary antibody. Development was performed using tyramide signal amplification. Hoechst 33342 (1:1000, Invitrogen) was used to stain nuclei. Samples were mounted using Scale A2 as described by Adler and colleagues (93) and stored at 4 degrees.

### Antibody Generation and Western Blotting

Rabbit anti-DDX24 antibody was generated against the peptide CSPKSLNADMQQKMRLKKLE specific to SMED-DDX24 and affinity-purified (Abgenex, Bhubaneswar India). Western blotting was performed as described by Owlarn and colleagues (94). DDX24 antibody was used at 1:1000 dilution. Rabbit Beta-Actin antibody (Cell Signaling Technology 4967S) and Rabbit SMED-PIWI1 antibody (60) were used at 1:2000 & 1:1000 dilution respectively.

### Image Acquisition

All brightfield images were taken Olympus SZX16. Confocal images were acquired on an Olympus FV 3000 laser scanning microscope. Images were processed using Fiji. (https://imagej.net/Fiji). All cell counting was performed manually.

### RNA extraction, cDNA synthesis, and qRT-PCR analysis

For any experiment, ∼10 animals were used to extract RNA using TRIzol™ reagent (Invitrogen). cDNA was synthesised using SuperScript™ III Reverse Transcriptase (Invitrogen) using random hexamer primers. Maxima SYBR Green/ROX qPCR Master Mix (2X) (ThermoFisher Scientific) was used for qRT-PCR experiments.

qPCR primer pairs (5’ to 3’) used are:

**18S rRNA**-GGCGGTATTTTGATGACTCTGG (F), GTGGATGTGGTAGCCGTTTCTC (R)
**28S rRNA**-GATGCTTCCTATGAGTCGGATTG (F), CGGCTTCACTCGTTTACCTCTAA (R)
**ddx24**- GGATTTTTGAAACCGACACA (F), CCCAAATGCTAATGTTTTGC (R)
**Actin**- GCTCCACTCAATCCAAAAGC (F), TCAAATCTCTACCGGCCAAG (R)

Each experiment was repeated three times independently. Actin was used for normalisation. Unpaired t-test was used for inferencing statistical significance. All graph was plotted on GraphPad Prism 6.

### Polysome profiling

3 DPA planarian tails were soaked in cycloheximide (CHX) for 2 hours at 100 µg/ml final concentration in 1X planaria media. After this, they were macerated in cold CMF (calcium-magnesium free media) to dissociate them into cells. The cells were then pelleted by spinning at 300g for 10 minutes in a swinging bucket. The supernatant was discarded and cells were resuspended in 400 µl of polysome lysis buffer (200mM KCL, 5mM MgCl_2_, 0.01% TritonX-100, 20mM Tris-Cl pH 7.5, 1X Protease Inhibitor Cocktail, 40U/mL RNAse inhibitor, and 100 µg/ml CHX). Cells were incubated in lysis buffer for 30 min on ice, vigorously pipetted and vortexed. This was then centrifuged at 13,000g for 30 minutes. The supernatant was collected and was loaded onto a sucrose gradient (15% to 45%) and centrifuged at 40,000 RPM for 2 to 2.5 hours at 4°C. The sucrose gradient was then loaded onto an ISCO Teledyne UA-6 Absorbance Detector. Sixty percent sucrose was pumped from below to push the sucrose gradient to the UV chamber. The sensitivity parameter of the machine was set at 0.5.

### Transcriptome analysis

RNA was extracted from 3 DPA tail fragments and transcriptome library was prepared using NEBNext® Ultra™ II Directional RNA Library Prep with Sample Purification Beads (catalogue no-E7765L) kit and sequenced in Illumina HiSeq 2500 machine. All the samples were sequenced (as single-end) in biological replicates. Approximately 28 to 31 million reads were sequenced for every sample. These reads were adapter trimmed using Trimmomatic (95) and mapped to rRNA and other contamination databases. Reads that did not align with these databases were taken for further analysis. We used reference-based transcriptome assembly algorithms Hisat2 v2.1.0 (96); Cufflinks v2.2.1 (97) and Cuffdiff v2.2.1 (98) to identify differentially expressed transcripts. We used Hisat2 (-q -p 8 --min-intronlen 50 --max-intronlen 250000 --dta-cufflinks --new-summary --summary-file) to align the reads back to dd_Smes_G4 (99) assembly of *Schmidtea mediterranea* genome. Around 81-84% of reads were mapped to dd_Smes_G4 genome. We used samtools to obtain sorted bam files. The mapped reads were assembled using Cufflinks (-p -o -b -u -N --total-hits-norm -G) with the most recent and well-annotated SMEST transcriptome as reference. We used cuffmerge to merge the gene list across different conditions. We identified differentially expressed genes using Cuffdiff module (-p –o ./ -b -u -N --total-hits-norm -L) and considered genes with adjusted p-value <0.05 as significance cut-off. Genes with significant p-value and at least two-fold up/down regulation were considered for further analysis. We did pathway analysis & gene-ontology analysis for these selected up/down regulated transcripts using GSEA (100, 101). We used a customized perl script for all the analyses used in this study. We used R ggplot2 (102), pHeatmap and CummeRbund (103) library for plotting. Different planarian cell-type markers were obtained from the available single-cell transcriptome data (61).

### Single-cell transcriptome analysis

We used single-cell transcriptome published from Peter Reddien’s lab (61) to extract the cells that express *ddx24*. We used the data matrix submitted in SRA to extract only the cells that express *ddx24*. We found 3929 cells (out of 50,563 cells) expressing *ddx24* transcript in the single-cell transcriptome. We re-analyzed the single-cell data as described in Ross and colleagues (104). We used Seurat (https://satijalab.org/seurat/), an R package to analyze the single-cell transcriptome for the cells that express *ddx24* mRNA (105, 106). Based on the markers from Fincher *et al.,* 2018, we classified the UMAP clusters as cell types. We used LogNormalize method of Seurat to normalize the dataset which was further scaled (linear transformation) using Seurat. This scaled value was further log-transformed and plotted as a heatmap for genes of interest. We used R ggplot2 GMD and heatmap.2 to derive all the plots.

### Fluorescence-activated cell sorting (FACS)

FACS was performed as described by Wagner and colleagues (56).

## ACKNOWLEDGMENT

We are grateful to Prof. Apurva Sarin for her critical comments and for proofreading our manuscript. SRS and VKD thank NCBS-inStem Ph.D. fellowship for funding. MMH is supported by DBT-JRF and VL is supported by CSIR-SRF. AJ and SS thank inStem for funding. We thank all the members of the Palakodeti and Sowdhamini labs for their constant support, critique, encouragement, and feedback, especially Dhiru Bansal, Giulia Moreni, Sayan Biswas, and Snehal Karpe for helping with experiments. We are grateful to Nishan Shettigar and Anirudh Chakravarthy from Akash Gulyani’s lab for sharing their unpublished anti- Gα q/11/14 antibody staining protocol (Shettigar et al., 2020, under review) and to Prof. Kiyokazu Agata for generously gifting us anti-arrestin antibody. Authors acknowledge the in-house central imaging and flow cytometry facility (CIFF) and NGS facility (especially Awadhesh Pandit) for constant technical support. The work was funded by NCBS core grants and JC Bose fellowship to RS (SB/S2/JC-071/2015) and DST Swarnajayanti fellowship (DST/SJF/LSA-02/2015-16) and inStem core grants awarded to DP.

## AUTHOR CONTRIBUTIONS

**Conceptualization**: Souradeep R. Sarkar, Dasaradhi Palakodeti.

**Data curation:** Souradeep R. Sarkar, Vairavan Laxmanan.

**Formal analysis:** Souradeep R. Sarkar, Vairavan Laxmanan.

**Funding acquisition**: Ramanathan Sowdhamini, Dasaradhi Palakodeti.

**Investigation:** Souradeep R. Sarkar, Vinay Kumar Dubey, Anusha Jahagirdar, Mohamed Mohamed Haroon, Sai Sowndarya.

**Methodology:** Souradeep R. Sarkar

**Project administration:** Souradeep R. Sarkar.

**Resources:** Ramanathan Sowdhamini, Dasaradhi Palakodeti.

**Software:** Vairavan Laxmanan.

**Supervision:** Ramanathan Sowdhamini, Dasaradhi Palakodeti.

**Validation:** Souradeep R. Sarkar, Anusha Jahagirdar, Mohamed Mohamed Haroon.

**Visualization:** Souradeep R. Sarkar.

**Writing – original draft:** Souradeep R. Sarkar.

**Writing – review & editing**: Souradeep R. Sarkar, Vairavan Laxmanan, Vinay Kumar Dubey, Anusha Jahagirdar, Mohamed Mohamed Haroon, Dasaradhi Palakodeti.

---

**Figure 1 (Supplementary).**
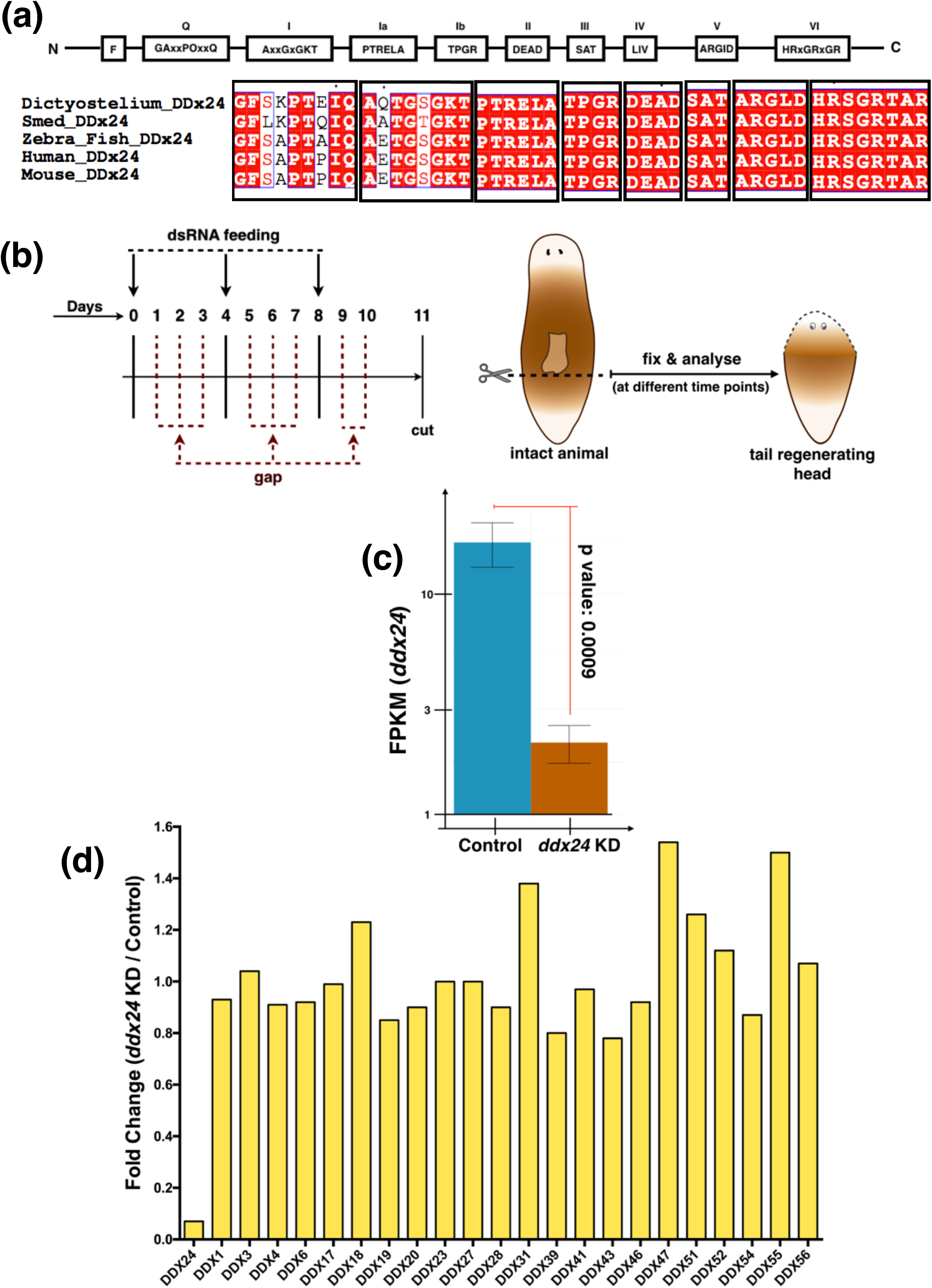
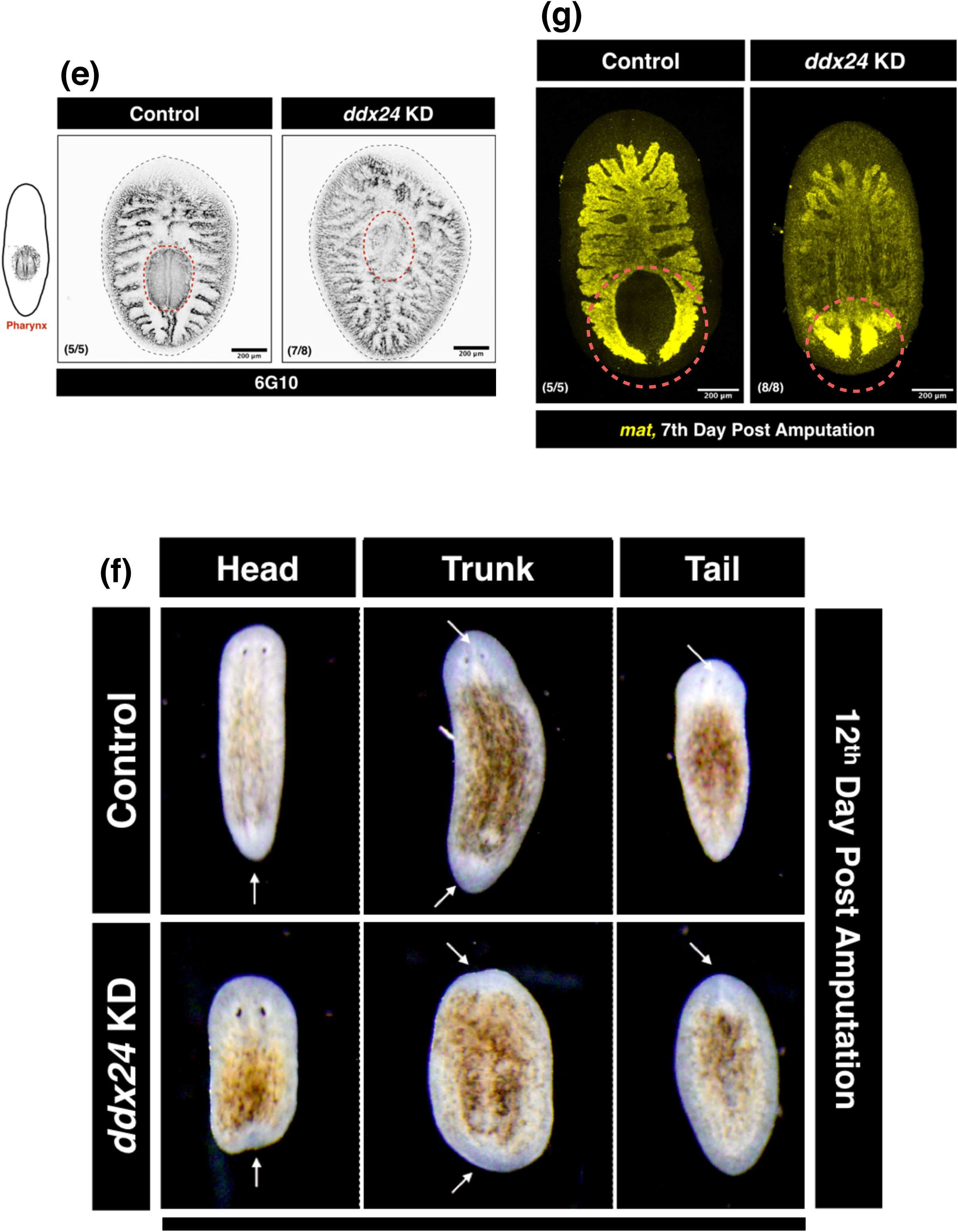
**(a)** Multiple sequence alignment for DDX24 from different species. We found SMED-DDX24, like DDX24 from other species, contains the motifs conserved across DEAD-box helicases. **(b)** Scheme and timeline of *ddx24* RNAi. **(c)** Efficiency of *ddx24* RNAi was determined by RNA-Seq. RNA was extracted on the 3rd day-post-amputation. **(d)** *ddx24* RNAi was specific. Even though planarians contain many DEAD Box Helicases, RNAi against *ddx24* was specific since transcript levels of *ddx24* alone got reduced. **(e)** Defective pharynx regeneration in *ddx24* KD animals. 6G10, which also marks pharyngeal muscle, was used to probe defective pharynx regeneration. (Scale bar: 200 µm) **(f)** Loss of DDX24 leads to an overall loss of regeneration in planarians. Head, trunk, and tail, all failed to regenerate eyespots and/or tails. For example, head fragments regenerating a tail failed to form the posterior gut branches **(g).**

**Figure 2 (Supplementary).**
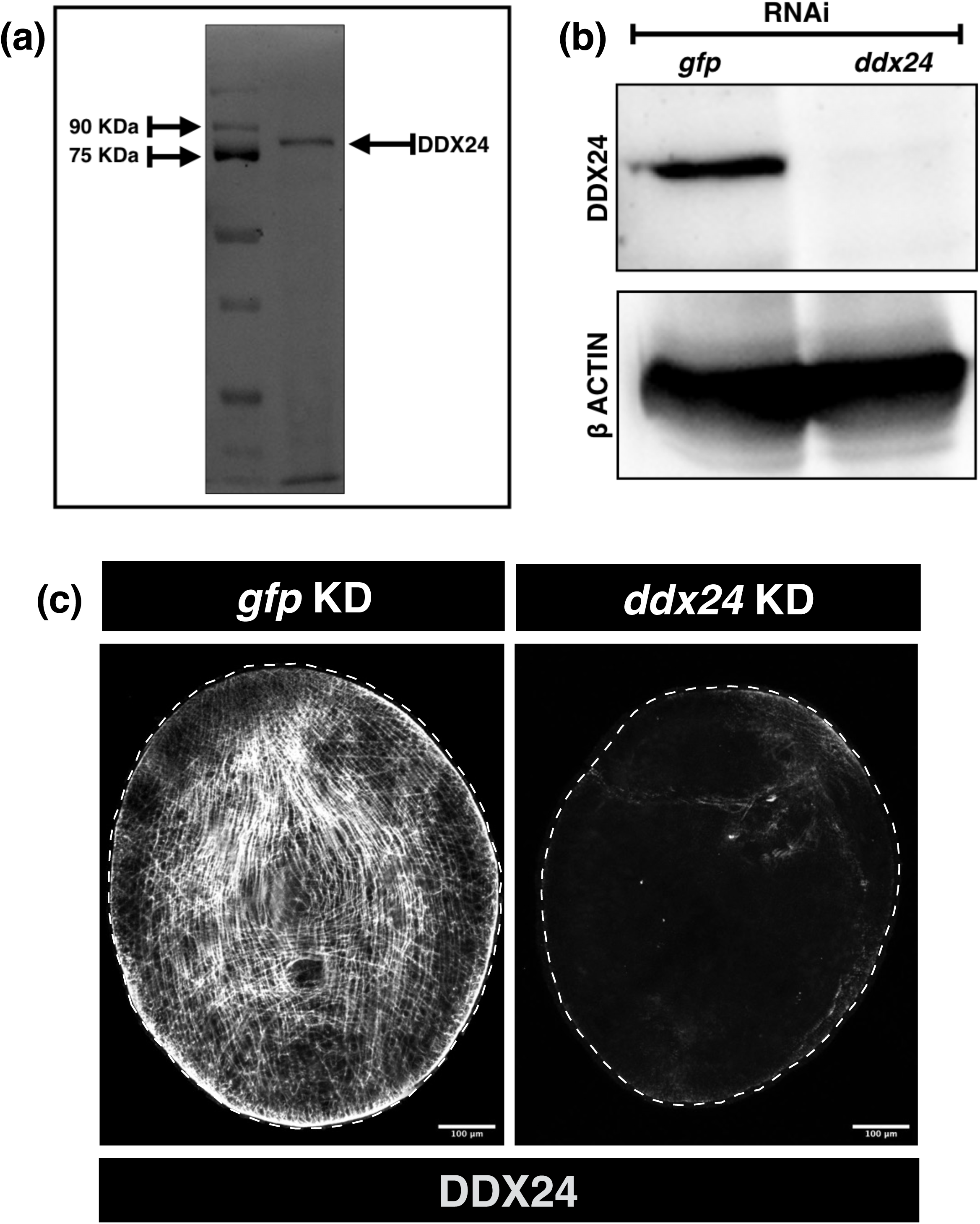
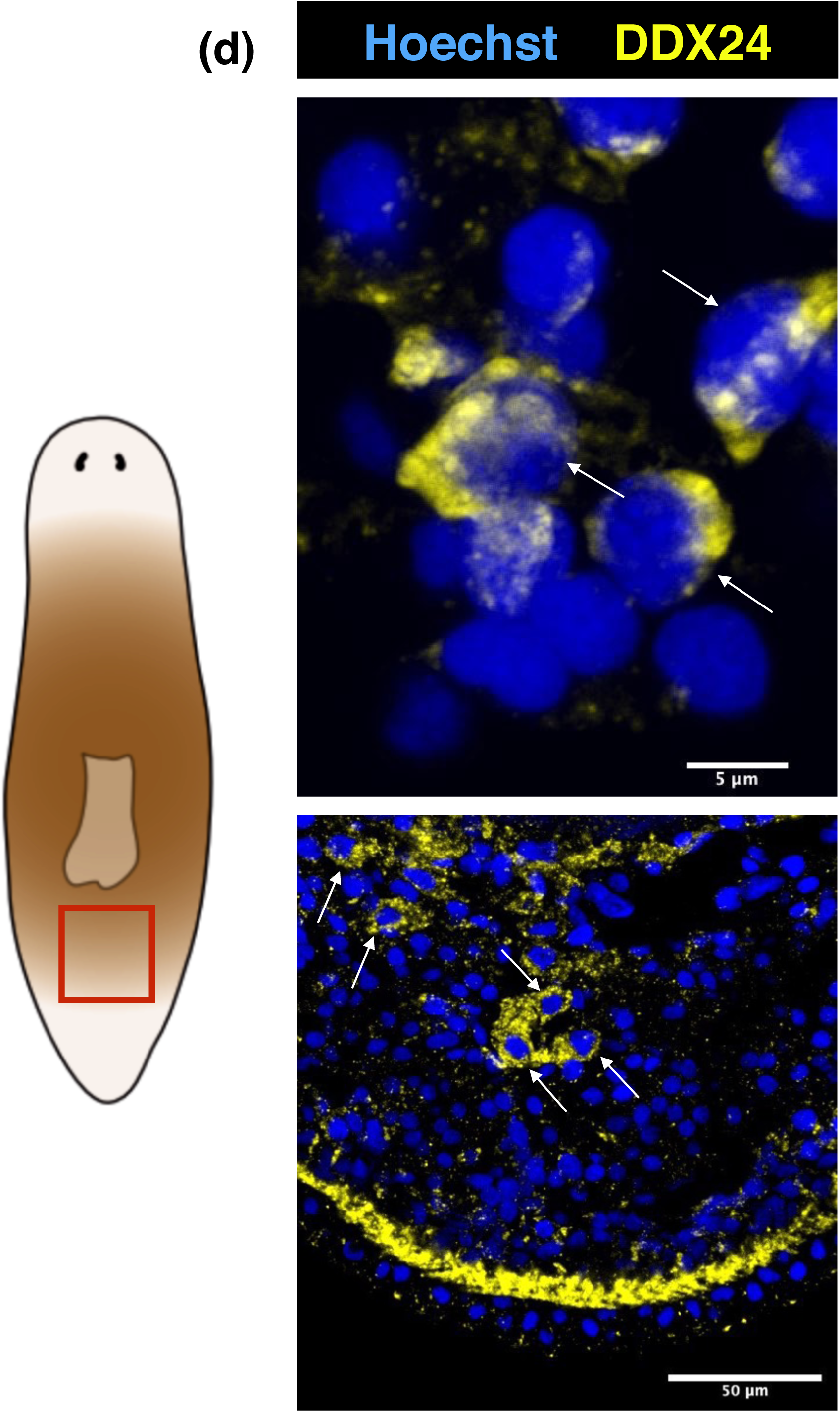
**(a)** Western blot using anti-DDX24 antibody gave a band above 75KDa but lower than 90KDa. **(b, c)** Validating the specificity of anti-DDX24 antibody by western blot ((b), N>5) and by immunostaining (c). **(d)** In addition to being enriched in muscle fibers, we found other DDX24 protein+ cells in the post-pharyngeal space. Since we were unable to perform DDX24 immunostaining post-RNA-FISH, we couldn’t confirm the identity of these cells.

**Figure 3 (Supplementary).**
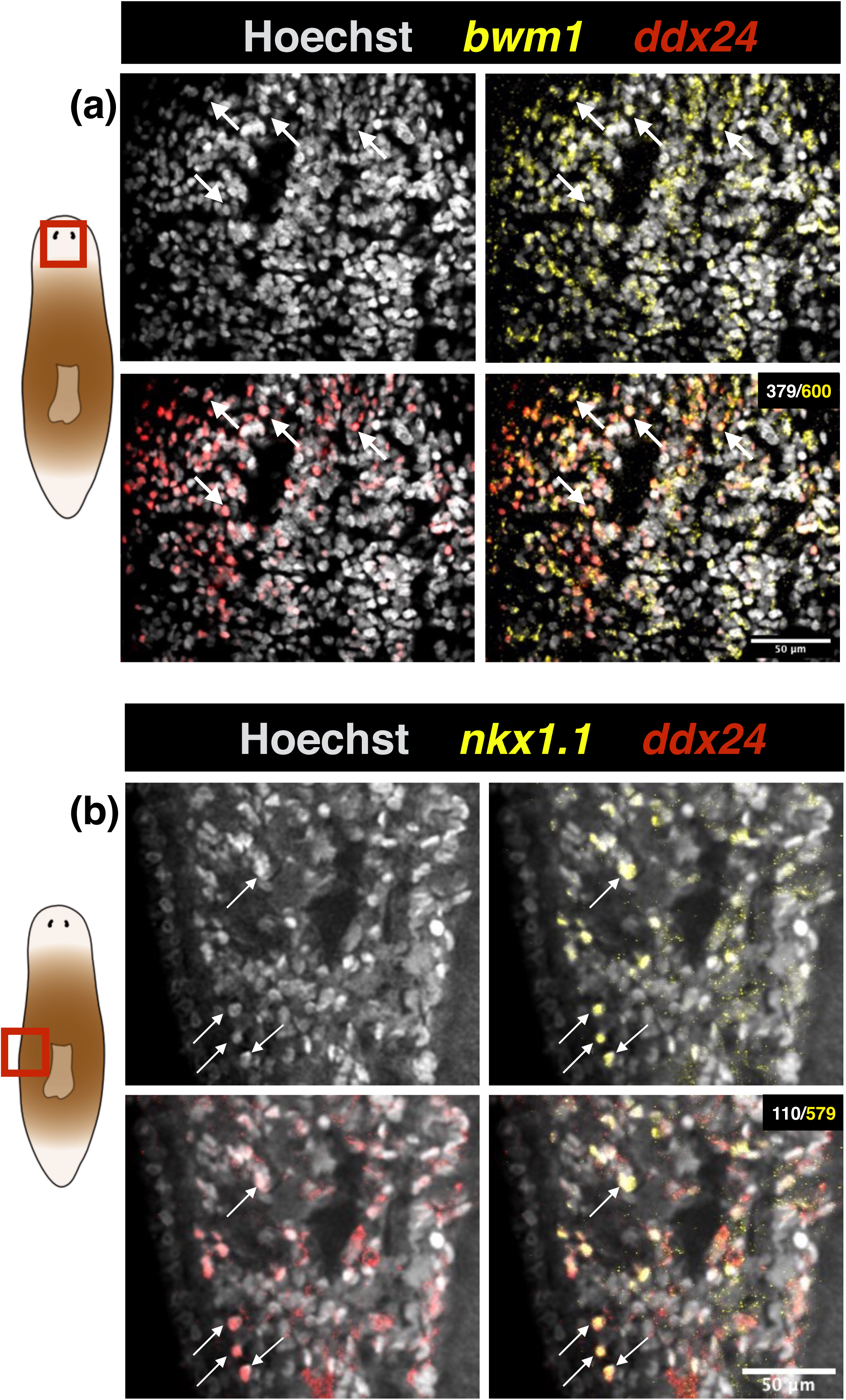
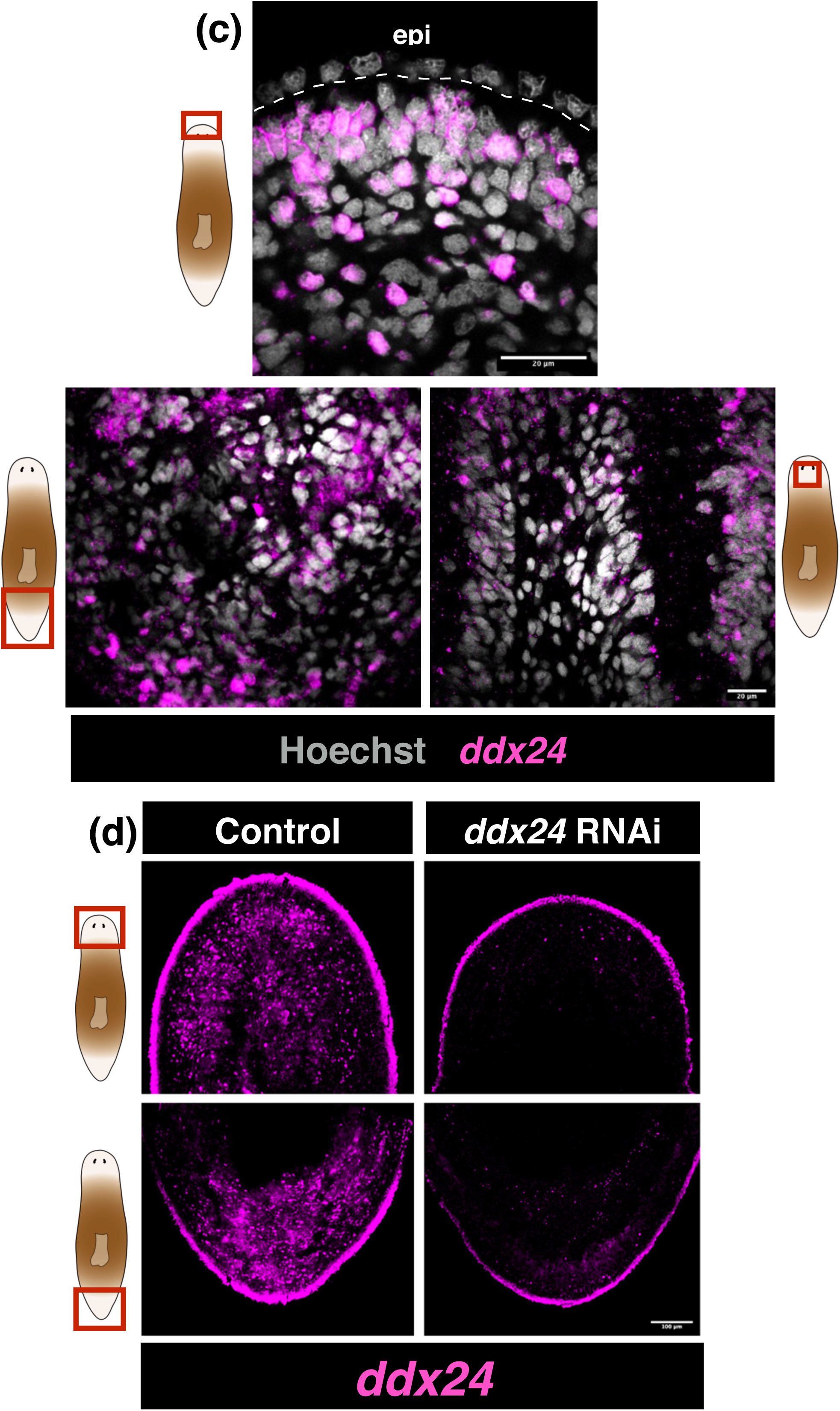
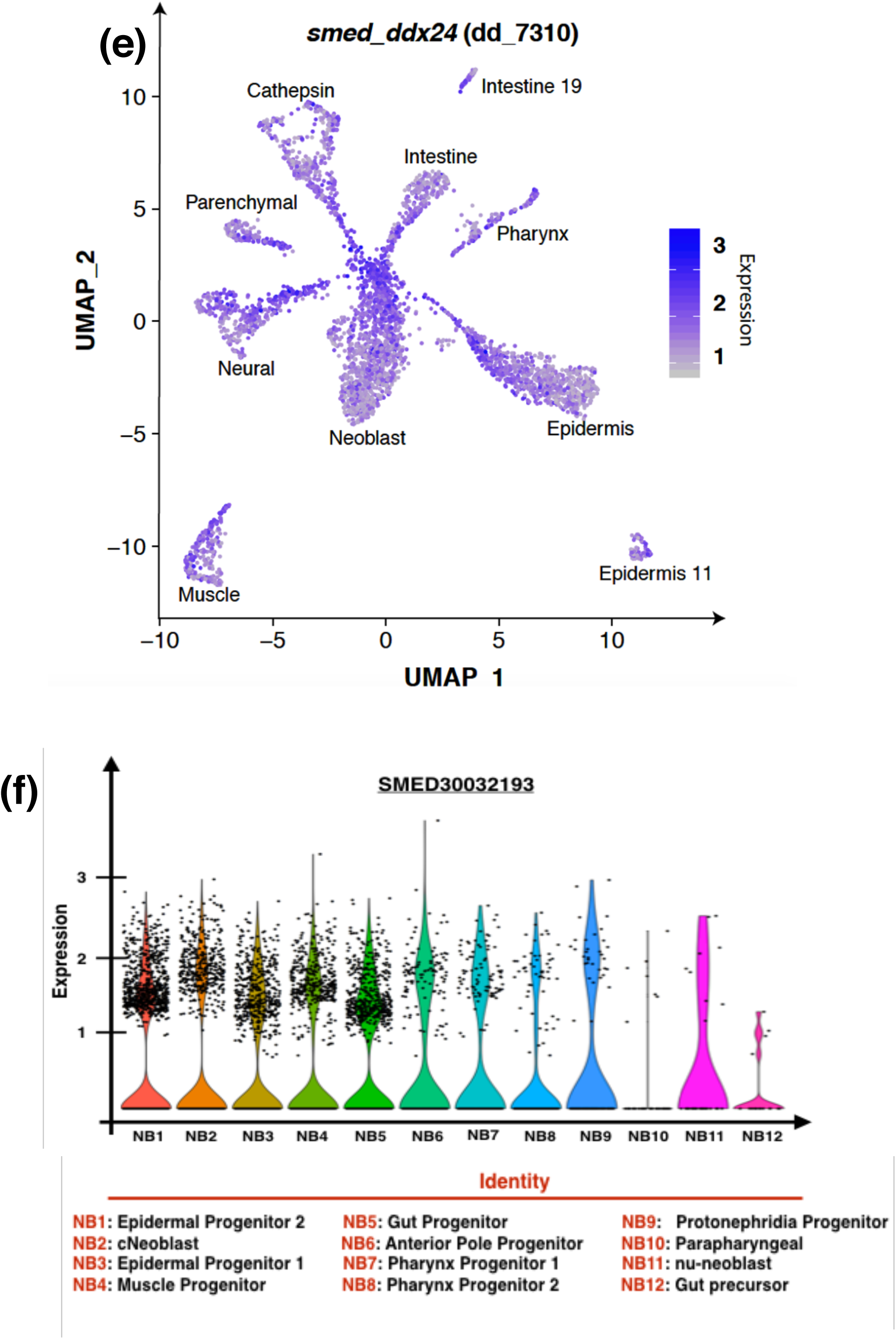
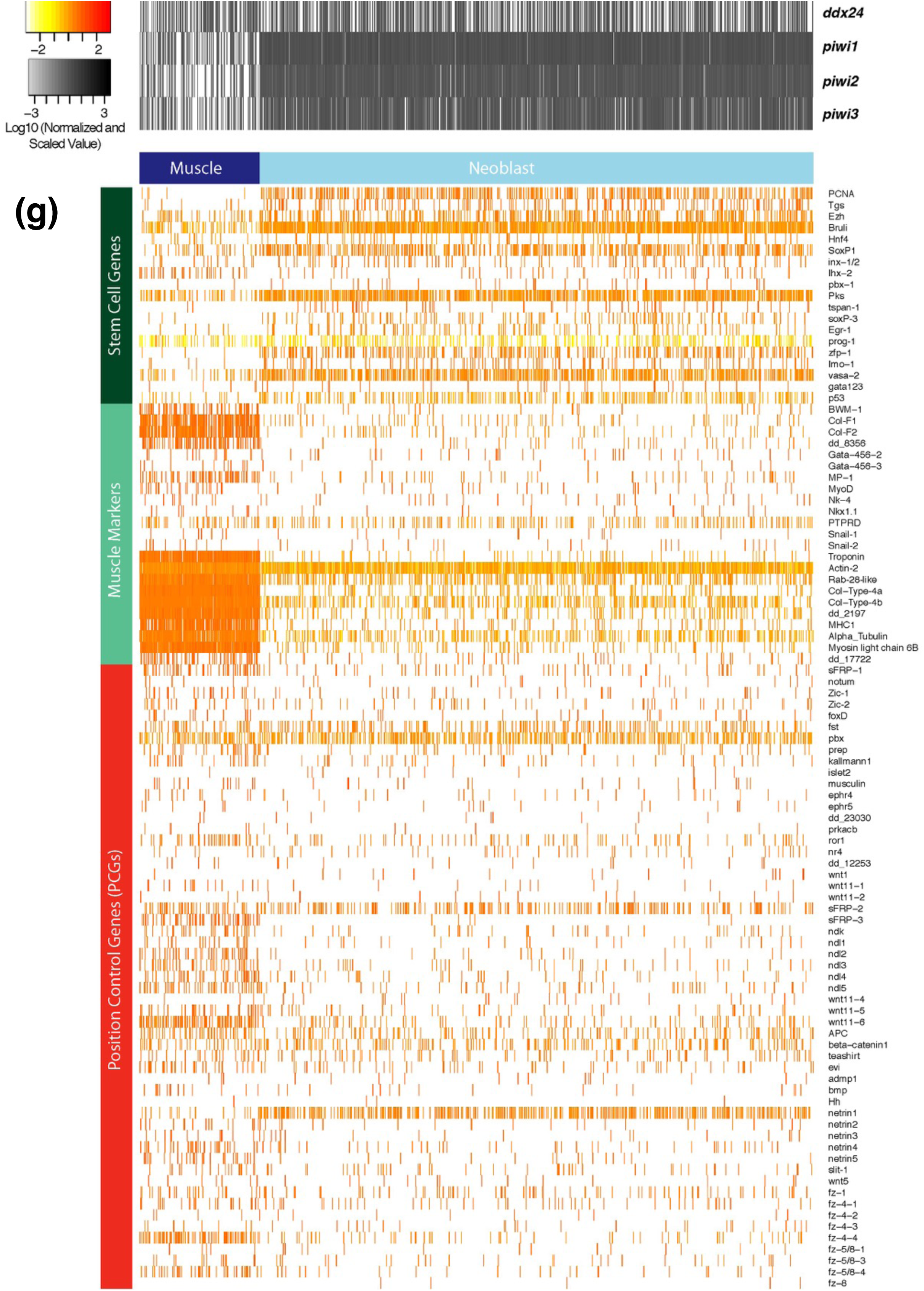
**(a, b)** *ddx24* mRNA also co-localized with *bwm1* and *nkx1.1*. In addition to *collagen* and *myoD*, these two are also bonafide muscle markers in planarians. 63.4±7.6% and 21.3±7.8% *bwm1+* and *nkx1.1+* cells respectively were also positive for *ddx24*. (Scale bar: 50 µm, 3 animals were used for quantification) **(c)** RNA-FISH for *ddx24*. Different regions highlighted-head tip, post-pharyngeal space, and cephalic ganglia. *ddx24* transcript was found to be enriched more on the ventral side of the animals as compared to the dorsal side. (scale bar: 20 µm) (**d)** *ddx24* RNA-FISH validation. (Scale bar: 100 µm) **(e)** Single-cell RNA sequencing profile for *ddx24* across different tissue types (Fincher et al., 2018). Supplementary table S4, Fincher et al., 2018, also listed *ddx24* as a posterior muscle enriched transcript. **(f)** Single-cell RNA sequencing profile for *ddx24* across different X1 neoblast clusters (Zeng et al., 2019). In addition to other lineages, both (e, f) suggested that *ddx24* was expressed in a subset of neoblasts and muscles, including specialized muscles are known as anterior-pole progenitor cells. **(g)** *ddx24+* single cells from Fincher et al., 2018, were extracted and re-analysed for the co-expression of other transcripts, as described by Ross et al., 2018. We overlaid the expression of selected stem cell genes, muscle genes, and positional-control and patterning genes on the *ddx24+* single cells. This co-expression dataset helped us corroborate the veracity of our RNA-FISH imaging data.

**Figure 4 (Supplementary).**
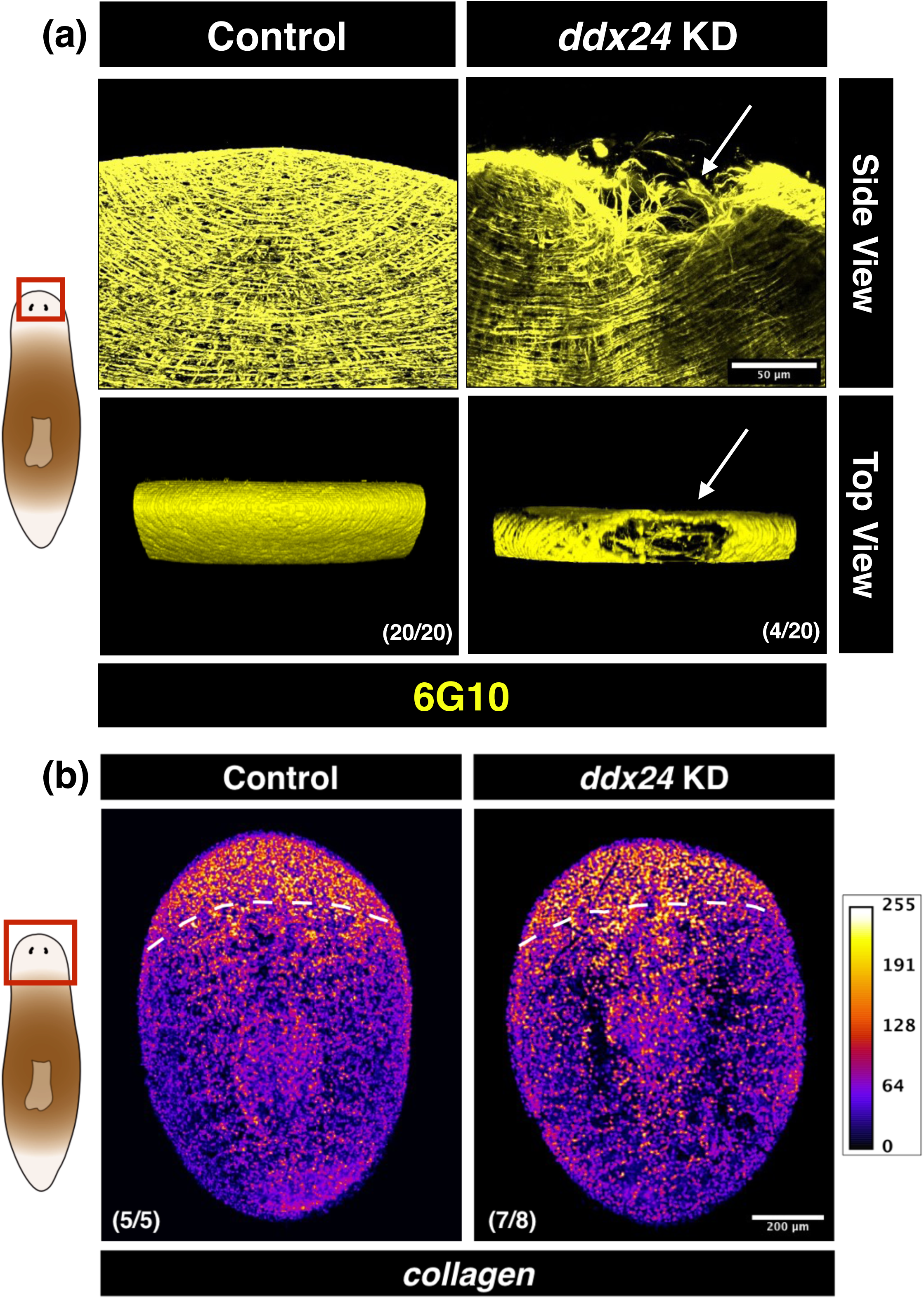
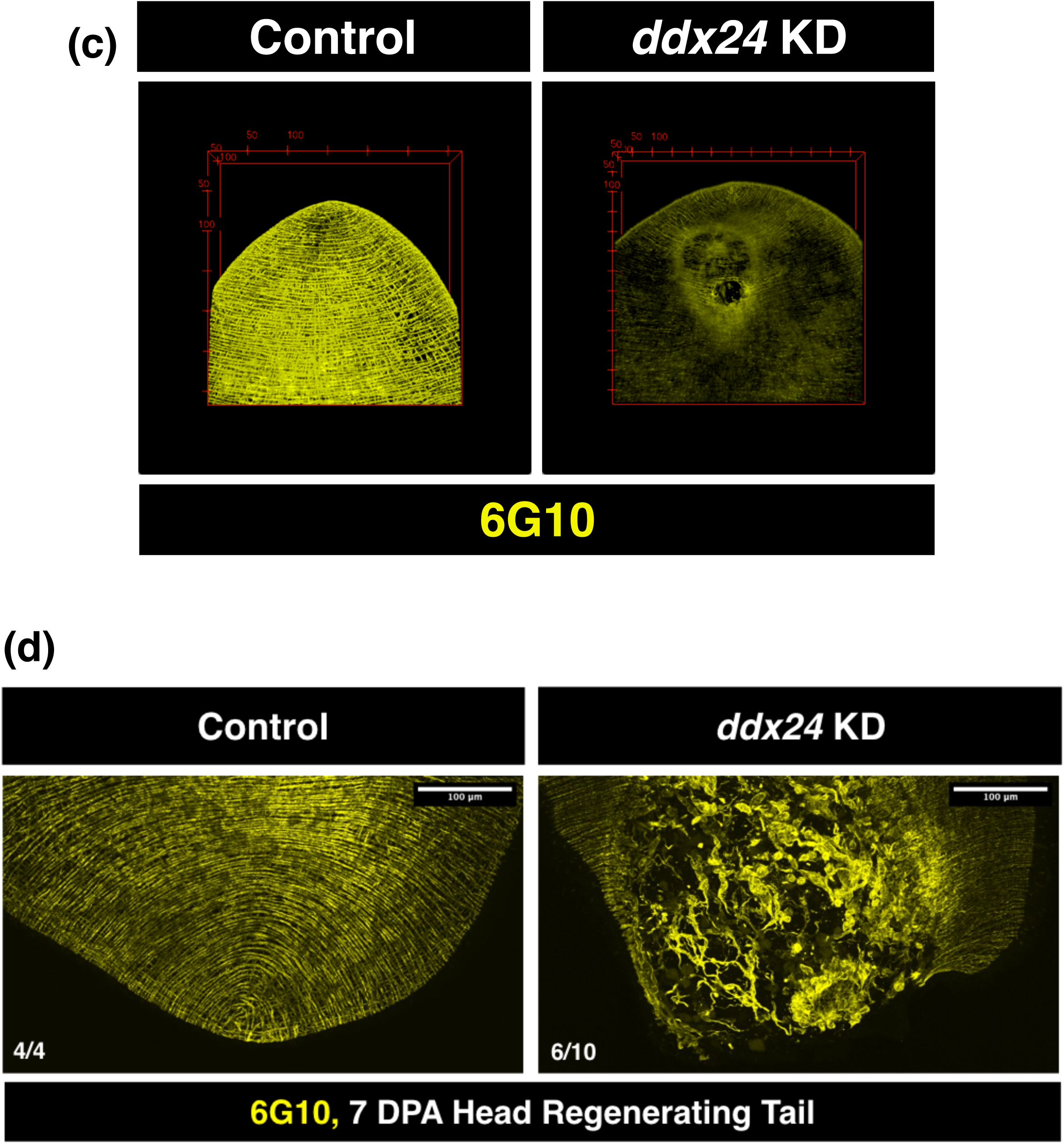
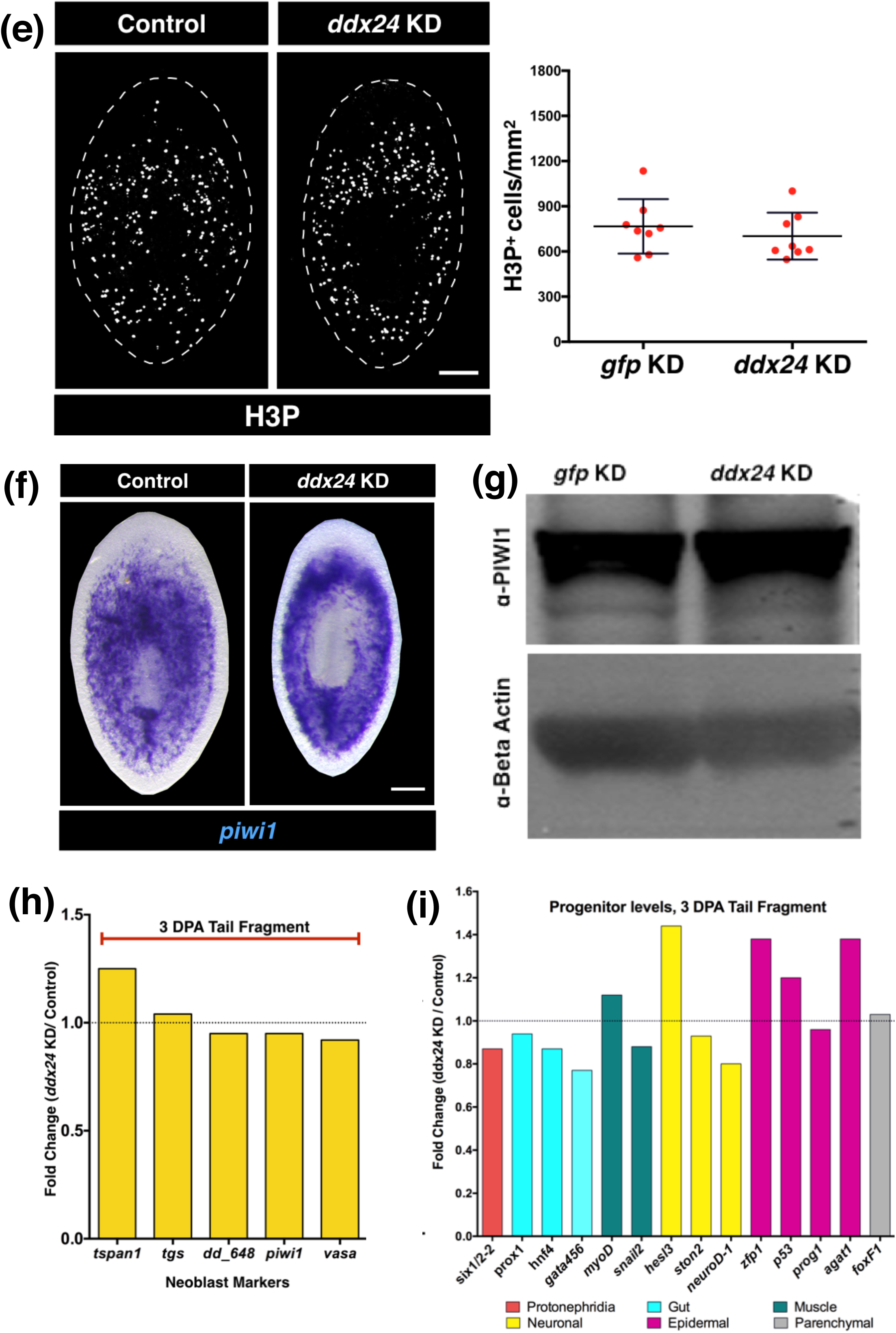
**(a)** Some *ddx24* KD animals displayed muscle fiber indentation at the anterior tip. This region houses a group of specialized muscle cells called anterior pole cells which are necessary for head regeneration as well as patterning of different anterior tissues. (Scale bar: 50 µm) **(b)** *collagen* mRNA is a bonafide muscle marker in planarians. RNA-FISH for *collagen* suggested that *ddx24* KD animals have similar collagen levels w.r.t. control fragments. This implied that neoblasts differentiated into *collagen+* muscle cells but the muscle fibers somehow failed to organize themselves appropriately in absence of DDX24. (Scale bar: 200 µm) **(c)** 3D movie showing 6G10 staining in control and *ddx24* KD animals. Muscle fiber rupture on both the dorsal as well as on the ventral surface can be seen here. Also, 6G10 stained *ddx24* KD animals weakly w.r.t control animals. All staining was performed on 7 DPA tail fragments. **(d)** Disruption of muscle fiber architecture was also observed in 7 DPA *ddx24* KD head fragments regenerating tail. (Scale bar: 100 µm) Although a subset of neoblasts was positive for DDX24 protein as well as for mRNA, loss of DDX24 did not affect neoblasts or their proliferation **(e, f, g, h)**. H3P makers proliferating neoblasts, *piwi1* mRNA is a pan-neoblast marker, whereas PIWI1 protein is present in all naïve neoblasts, primed neoblasts as well as their immediate early progeny. In addition, Log2FoldChange values for different neoblast associated transcripts (*tspan1*, *tgs1*, *dd_648*, *vasa1*, and *piwi1*) from RNA sequencing performed on 72 hpa tails suggested that their levels were comparable in *ddx24* KD animals w.r.t. controls. **(j)** Log2FoldChange values for different progenitor transcripts indicated that DDX24 KD animals did not display differentiation defect. RNA-Seq was performed in 3 DPA tail fragments. This data was further corroborated by RNA-FISH for collagen on 7 DPA animals. collagen differentiation from *piwi1+* neoblasts seemed unperturbed in *ddx24* KD animals (Figure 4-figure supplement 4b).

**Figure 5 (Supplementary).**
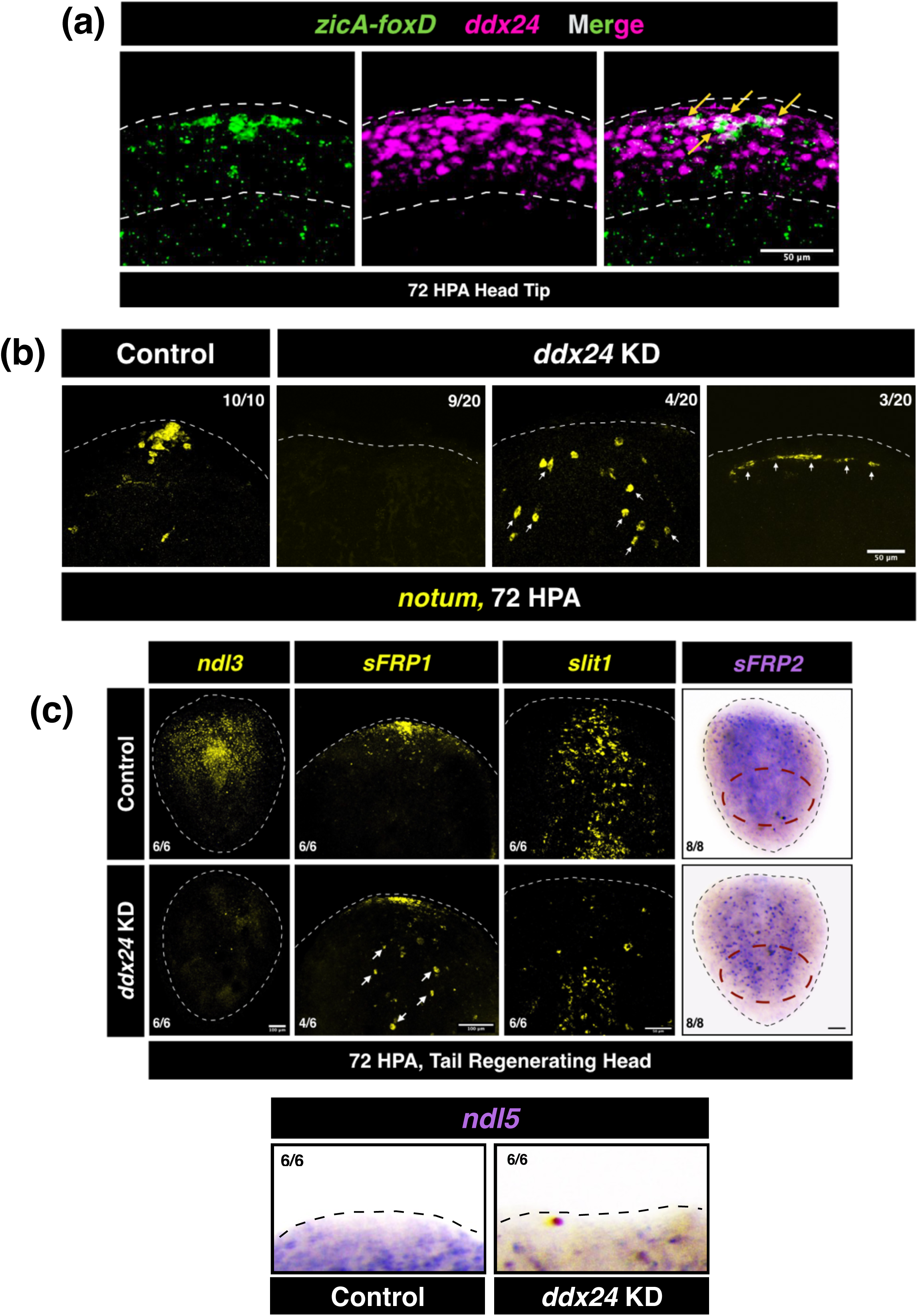
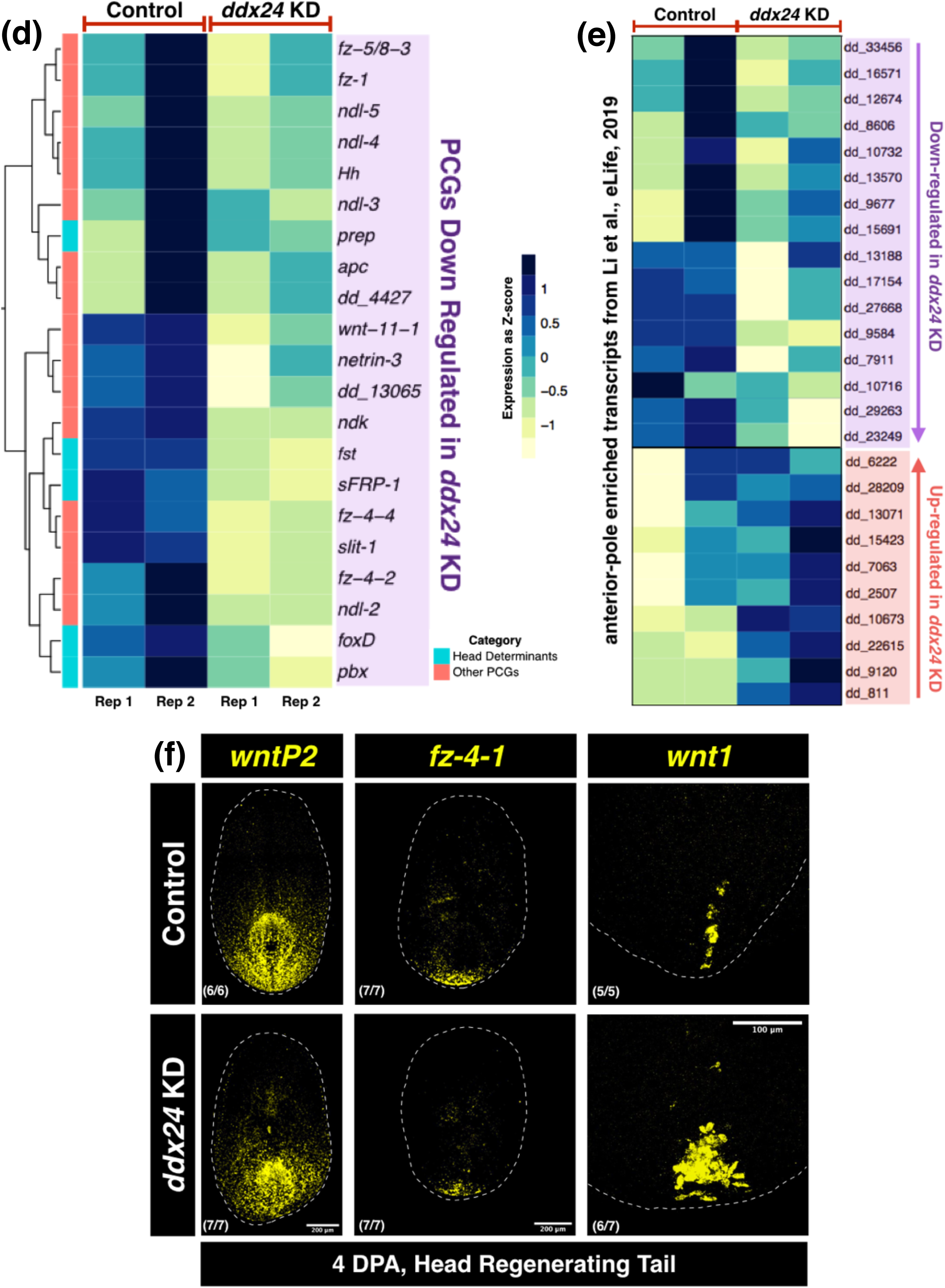
**(a)** *ddx24* was expressed at the 72 hpa head blastema region (area enclosed by white dashed lines). During this time, *ddx24* was also expressed by *zicA+foxD+* anterior pole progenitor cells in this region (yellow arrows) (Scale bar: 50 µm). **(b)** *notum* expression was also reduced at the pole region in a subset of *ddx24* KD animals by 72 hpa. In other animals, mis-expression of *notum* (white arrows) was observed. (Scale bar: 50 µm). **(c)** Loss of DDX24 leads to a reduction in expression and/or mis-expression of different position-control genes and patterning genes required for bonafide head identity. **(d)** We performed bulk RNA sequencing from 3DPA (72HPA) tail fragments regenerating the head. In addition to corroborating our staining above, we saw that expression of many other well-known genes essential for proper head regeneration and patterning were down-regulated in *ddx24* KD animals. **(e)** Dayan Li et al., 2019, provided a comprehensive list of transcripts up-regulated in the head primordia by 3DPA (72 HPA). Comparing our transcriptome data with this paper gave us two sets of genes- (1) one set was down-regulated in *ddx24* KD and (2) the other set was up-regulated in *ddx24* KD. This further seemed to suggest that anterior pole determination was abnormal in absence of DDX24. **(f)** PCG expression during tail regeneration in *ddx24* KD animals. Although expression of tail determinants/markers like *wntP2/wnt11-5* and *fz-4-1* weren’t aberrant in absence of DDX24, the number of *wnt1+* cells at the tail tip was highly upregulated.

